# Olfactory and vomeronasal innervation of the olfactory bulbs are not essential for GnRH-1 neuronal migration to the brain

**DOI:** 10.1101/154922

**Authors:** Ed Zandro Taroc, Aparna Prasad, Jennifer M. Lin, Paolo E. Forni

**Affiliations:** Department of Biological Sciences, University at Albany, Albany, NY, 12222

## Abstract

Gonadotropin releasing hormone-1 (GnRH-1) neurons migrate from the developing olfactory pit into the hypothalamus during embryonic development. Migration of the GnRH-1 neurons is required for mammalian reproduction as these cells control release of gonadotropins from the anterior pituitary gland. Disturbances in GnRH-1 cell migration, GnRH-1 synthesis, secretion or signaling lead to varying degrees of hypogonadotropic hypogonadism (HH), which impairs pubertal onset and fertility. HH associated with congenital olfactory defects is clinically defined as Kallmann Syndrome (KS).

The association of olfactory defects with HH in KS suggested potential direct relationship between defective olfactory axonal routing, lack of olfactory bulbs and aberrant GnRH-1 cell migration. However, it has never been experimentally proven that the formation of axonal connections of the olfactory and vomeronasal neurons to their functional targets are necessary for the migration GnRH-1 neurons to the hypothalamus. Loss-of-function of the *Arx-1* homeobox gene leads to the lack of proper formation of the olfactory bulbs with abnormal axonal termination of olfactory sensory neurons (Yoshihara et al., 2005). We exploited the Arx-1^null^ mouse line to investigate the role of the olfactory system (olfactory/vomeronasal fibers and OBs) in controlling GnRH-1 migration to the hypothalamus. Our data proves that correct development of the OBs, and axonal connection of the olfactory and vomeronasal sensory neurons to the forebrain are not needed for GnRH-1 neuronal migration. Moreover, we prove that the terminal nerve, which forms the GnRH-1 migratory scaffold, follows different guidance cues and differs in its genetic expression from olfactory and vomeronasal sensory neurons.

**Significance Statement:** Gonadotropin Releasing Hormone-1 (GnRH-1) neurons control the reproductive axis of vertebrates. During embryonic development, these neurons migrate from the olfactory pit to the hypothalamus. GnRH-1 cell migration is commonly believed to take place along the olfactory axons. Our work reveals that correct olfactory bulb development and targeting of the olfactory and vomeronasal sensory neurons to the brain are not required for this migration. Our work challenges the idea that GnRH-1 neuronal migration to the hypothalamus relies on correct routing of the olfactory and vomeronasal sensory neurons. We provide a new basis for interpreting genetic correlations between anosmia, lack of olfactory bulbs, and hypogonadotropic hypogonadism in Kallmann Syndrome.

## Introduction

Gonadotropin Releasing Hormone I neurons (GnRH-1ns) play a pivotal role in controlling the reproductive axis of vertebrates. In the adult, the GnRH-1ns reside within the preoptic hypothalamic area (POA), where they control the hypothalamic–pituitary–gonadal hormonal axis (HPG axis), driving reproductive development and regulating reproductive hormones in adult life (Cattanach et al., 1977). During embryonic development, the GnRH-1ns originate in the developing olfactory pit (OP), from which they migrate into the brain and eventually arrive at the hypothalamus (Schwanzel-Fukuda and Pfaff, 1989; Wray et al., 1989a; Wray et al., 1989b). Disturbances either in this migration or in GnRH-1 synthesis, secretion, and signaling lead to hypogonadotropic hypogonadism (HH), which adversely affects normal sexual development, social interactions, fertility, and propagation of the species (Schwanzel-Fukuda et al., 1989; Burmeister et al., 2005; Yin and Gore, 2006; Maruska and Fernald, 2011; Zhang et al., 2013). Around half of all patients affected by HH either have difficulty perceiving odors (hyposmia) or entirely lack the ability to smell altogether (anosmia) (Bianco and Kaiser, 2009; Mitchell et al., 2011). HH associated with congenital olfactory defects is clinically defined as Kallmann Syndrome (KS).

Developing GnRH-1ns migrate along bundles of olfactory/vomeronasal (VN) and terminal nerve (TN) axons, which project from the nose to the olfactory bulb and POA, respectively. Whether these axons are collectively permissive for migration from the pit to the brain, or whether a specific subpopulation provides instructive guidance cues essential for successful migration has long been controversial. The prevailing idea is that GnRH-1ns access the brain along the olfactory/vomeronasal (VN) sensory fibers, primarily because HH in KS is generally associated with anosmia (Wray, 2010; Boehm et al., 2015; Forni and Wray, 2015). Studies based on genetically modified animal models have described GnRH-1ns migratory defects associated with an array of olfactory and vomeronasal sensory neuronal routing deficiencies, either alone or together with atypical formation of the olfactory bulbs (OBs)(Cariboni et al., 2007; Bergman et al., 2010; Cariboni et al., 2011; Cariboni et al., 2012; Hanchate et al., 2012; Messina and Giacobini, 2013; Pingault et al., 2013; Boehm et al., 2015; Cariboni et al., 2015; Lettieri et al., 2016).

Unilateral and bilateral absence or reduction in the size of OBs, are common phenotypes in Kallmann patients carrying Kal1, CHARGE, trisomy 13 or trisomy 18 mutations, Prok2 or Prok-R2 mutations, and mutations affecting Fgf8 signaling (Ng et al., 2005; Pitteloud et al., 2007; Hardelin and Dode, 2008; Dode and Hardelin, 2010; Teixeira et al., 2010). Defective projections to the CNS and defective bulb formation have also been described after loss of function of *Dlx5, Fezf1, Klf7, Emx2, Lhx2,* genes in mouse (Yoshida et al., 1997; Levi et al., 2003; Long et al., 2003; Yoshihara et al., 2005; Hirata et al., 2006; Chung et al., 2008; Berghard et al., 2012).

Despite many correlations between olfactory defects and aberrant GnRH-1ns migration, direct experimental evidence proving that olfactory and vomeronasal connections to the OBs are necessary for GnRH-1ns migration to the hypothalamus is lacking. Additionally, in families carrying mutations linked to KS, the two aberrant phenotypes, HH and anosmia, do not necessarily co-segregate.(Leopold et al., 1992; Yousem et al., 1996; Ghadami et al., 2004; Pitteloud et al., 2006; Frasnelli et al., 2007; Balasubramanian et al., 2014).

*Arx-1 is* an X linked homeobox gene related to the *Drosophila aristaless*. Arx-1 loss-of-function leads to a severe form of olfactory bulb aplasia together with abnormal axonal termination of olfactory sensory neurons (Yoshihara et al., 2005). We have exploited Arx-1^null^ mutants together with a series of reporter mouse models to test the role of the olfactory system in controlling GnRH-1 migration to the hypothalamus. Our data proves that the establishment of axonal connections of the olfactory and vomeronasal sensory neurons to the brain are not needed for GnRH-1 neuronal migration. In fact, the GnRH-1ns and terminal nerve appear to follow different guidance cues from those controlling the olfactory projections to the OBs.

## Material and Methods

### Animals

Cryopreserved Arx-1^null^ mice were resuscitated from the Riken repository. The origins of these mice and their olfactory defects were previously described (Kitamura et al., 2002; Yoshihara et al., 2005), and our Arx-1 colony was C57 BL/6J mixed background. Because the *Arx-1* gene is located on the X-chromosome, *Arx-1* hemizygous null mutants were also genotyped for sex to identify male mutants (see below). Arx-1^null^ mice do not survive past post-natal day 0 (P0), therefore animal cages were checked early in the morning on the day of birth. Embryos of different stages were collected after euthanizing time-mated dams. Peripherin-EGFP (hPRPH1-G) mice were obtained from Dr. J. Sasero (Murdoch Childrens Research Institute, Melbourne Australia) on a C57BL/6J background (McLenachan et al., 2008). hPRPH1-G Males were mated with Arx-1 females to generate hPRPH1-G /Arx-1^null^ embryos. GPR12EGFP BAC transgenic mice were resuscitated from GENSAT repository at MMRRC, UC Davis. These mice were obtained on a mixed background. Because EGFP expression in the VNO varied among the F_0-_resuscitated animals obtained from GENSAT, we selected a subline (GPR12EGFP^395^) with persistent strong expression in apical and basal vomeronasal sensory neurons. GPR12EGFP^395^ male mice were mated with Arx-1 females to generate Arx-1^null^/GPR12EGFP embryos. Animals were euthanized using CO_2_, followed by cervical dislocation. All animal procedures were in accordance with procedures approved by the University at Albany Institutional Animal Care and Use Committee (IACUC).

### Tissue preparation

Embryos were collected from time-mated *Arx-1* females, where the emergence of the copulation plug was taken as E0.5. Collected embryos were immersion-fixed in 3.7% Formaldehyde/PBS at 4°C for 3 hours. P0 heads were immersion-fixed in in the same fixative at 4°C for overnight. All samples were then cryoprotected in 30% sucrose overnight or until they sank, then frozen in O.C.T (Tissue-TeK) using dry ice, and kept at -80C. Samples were cryosectioned using CM3050S Leica cryostat and collected on Superfrost plus slides (VWR) at 12-16*μ*m for immunostainings and 18-25 *μ*m for *In Situ Hybridizations (ISH)*.

### Confirmation of animal genotypes

The genotypes of the mice were established by PCR analysis using the following primers: *Arx-1^null^ (*mArx fff: 5’-CGCCC AAGGA AGAGC TGTTG CTGC-3’; *ARX pMC1neo stop*: 5’-GCCTT CTTGA CGAGT TCTTC-3’; ARX mArx eer 5’TATTC CACCC TCCTG GACCC TTTC-3’); *EGFP (*eGFP fwd: 5’-CCTAC GGCGT GCAGT GCTTC AGC-3’; *eGFP REV:* 5’CGGCG AGCTG CACGC TGCGT CCTC-3’); *Actin 470 (ACTIN SENSE):* 5’-CTCGT CTGGGA AAGCA GAAAC TGCAA-3’; *ACTIN 470 (ANTISENSE)*: 5’-GTGAC CTGTT ACTGG GAGTG GCAAG C-3’). Amplification products were analyzed by agarose gel electrophoresis.

### Immunohistochemistry

Primary antibodies and dilutions used in this study were: goat-*α*-neuropilin-1(1:400, R&D Systems), goat-*α*-neuorpilin-2(1:3000, R&D Systems), goat-*α*-ROBO1 (1:200, R&D Systems), goat-*α*-ROBO3(1:50, R&D Systems), mouse-*α*-ROBO2(1:50, Santa Cruz) chicken-*α*-peripherin (1:1500, Abcam), rabbit-*α*-peripherin (1:2000, Millipore), SW rabbit-*α*-GnRH-1 (1:6000, Susan Wray, NIH), rabbit-*α* tyrosine hydroxylase (1:1000, Abcam), goat-*α* olfactory marker protein (1:4000, WAKO), goat-*α* transient-axonal glycoprotein 1(1:1000, R&D Systems), rabbit-*α*-GFP(1:2000, Molecular Probes), chicken-*α*-GFP (1:1000, Abcam), mouse -*α*-GAD67 (1:200, Santa Cruz). Antigen retrieval was performed in a citric acid solution prior to incubation with chicken-*α*-peripherin, rabbit-*α*-tyrosine hydroxylase, mouse-*α*-ROBO2, and *α*-GAD67 antibodies. For immunoperoxidase staining procedures, slides were processed using standard protocols (Forni et al., 2013) and staining was visualized (Vectastain ABC Kit, Vector) using diaminobenzidine (DAB) in a glucose solution containing glucose oxidase to generate hydrogen peroxide; sections were counterstained with methyl green. For immunofluorescence, species-appropriate secondary antibodies were conjugated with Alexa-488, Alexa-594, or Alexa-568 (Molecular Probes and Jackson Laboratories) as specified in the legends. Sections were counterstained with 4′,6′-diamidino-2-phenylindole (1:3000; Sigma-Aldrich) and coverslips were mounted with Fluoro Gel (Electron Microscopy Services). Confocal microscopy pictures were taken on a Zeiss LSM 710 microscope. Epifluorescence pictures were taken on a Leica DM4000 B LED fluorescence microscope equipped with a Leica DFC310 FX camera. Images were further analyzed using FIJ/ImageJ software.

### X-gal Staining

Sections were rehydrated in PBS and then incubated in a solution of 5 mM potassium ferrocyanide, 5 mM potassium ferricyanide, 2 mM MgCl_2_, 0.1% Tween, and 0.1% 5-bromo-4-chloro-3-indolyl-b-D-galactoside/dimethylformamide at 37°C O.N. After the enzymatic reaction was completed, slides were either counterstained with 1% Eosin-Y (Electron Microscopy Services) or washed and immunostained as described above.

### In situ hybridization

Digoxigenin-labeled cRNA probes were prepared by *in vitro* transcription (DIG RNA labeling kit; Roche Diagnostics) from the following templates: Semaphorins 3A, (Kagoshima and Ito, 2001); Slits-1, as well as Robo-2, Robo-3 (Cloutier et al., 2004). In situ hybridization was performed as described (Forni et al., 2013) and visualized by immunostaining with an alkaline phosphatase conjugated anti-DIG (1:1000), and NBT/BCIP developer solution (Roche Diagnostics). Sections were then counter-immunostained with antibodies against both chicken-*α*-peripherin, and SW rabbit-*α*-GnRH-1, as described above for immunofluorescence.

### Quantification and statistical analyses of microscopy data

Cell counts were done on serial sections immunostained for GnRH-1 and visualized under bright field (immunoperoxidase) or epi-fluorescence illumination (20×; Leica DM4000 B LED), according to their anatomical location [*i.e.,* (1) nasal region (VNO, axonal tracks surrounding the olfactory pits, forebrain junction); (2) olfactory bulb/fibrocellular mass; and (3) brain (all the cells that accessed the olfactory bulb and were distributed within the forebrain)]. For each animal, counts were performed on 3 serial series. The average number of cells from these 3 series was then multiplied by the total number of series/animal to compute a value for each animal. These were then averaged (± standard error) among animals of the same age and genotype. Means ± SEs were calculated on at least three animals per genotype. A two-tailed, non-paired Student’s *t*-test was used to assess differences between mutant and wild type controls for each group. P≤ 0.05 was considered statistically significant.

### Mapping the Distribution of GnRH-1 Neurons

Whole heads of P0 Arx-1^null^ mutants and controls (n=3;3) were cryosectioned at 16*μ*m thickness. The sections were then immunostained against GnRH-1 in DAB and counterstained with methyl green. The most medial 16 sections (8 sections from either side of the midline cartilage) were scanned at 10x using a VS120 Olympus scanning microscope in brightfield. Sections were aligned in PhotoShop CS6 using the median eminence, cerebellum, and ventricles as landmarks. Cell bodies were marked and overlaid, representing a cross section of their migratory path. The coordinates of each cell body were plotted in reference to the origin (x=0;y=0), which was set at the middle of the median eminence, using FIJI. The number of GnRH-1ns distributed along the x-axis (rostro-caudal) were quantified in 500*μ*m intervals for each animal. Differences at each interval between genotypes was assessed by unpaired t-Test.

## Results

### Arx-1 mutants lack proper olfactory bulb formation

As previously described in detail by Yoshihara and coworkers (Yoshihara et al., 2005), Arx-1^null^ mice develop a severe bulb aplasia/hypoplasia secondary to the defective proliferation, migration, and maturation of interneuron progenitors and precursors into the OB. Periglomerular cells and granule cells are two major types of GABAergic interneurons in the OB (Mugnaini et al., 1984; Kosaka et al., 1995; Kiyokage et al., 2017). Tyrosine hydroxylase (TH) is expressed by sets of periglomerular cells and cells of the molecular layer. Olfactory nerve input is required for the normal expression of tyrosine hydroxylase (TH) in the main olfactory bulb (Kawano and Margolis, 1982; Ehrlich et al., 1990; Stone et al., 1990).

Control mice immunostained for olfactory marker protein (OMP; to label olfactory and vomeronasal neurons and axons) and TH revealed the normal projections and active connections of olfactory/vomeronasal axons (Fig. 1A,C,I,K). In Arx-1null animals, the olfactory fibers were found tangled in a large fibro-cellular mass (FCM; Fig. 1B,D,J,L) and no TH immunoreactivity was found in the OBs (Fig. 1J,L).

Immunostaining against glutamic acid decarboxylase-67 (GAD67) (Mugnaini et al., 1984; Carleton et al., 2003), highlighted the well-organized GABAergic neurons in the OB of control animals (Fig. 1M). In Arx-1 mutants most of the GAD67 positive neurons could not enter the OB and accumulated ventral and at the rostral end of the RMS (Fig. 1N). However, comparable TH and GAD67 immunoreactivity was found in the striatum of WT and Arx-1^null^ mutants (Fig 1I,J,M,N, asterisks).

In the Arx-1^null^ mice, two exons have been replaced by the β-galactosidase gene (Kitamura et al., 2002). By performing X-Gal staining on Arx-1^null^ mutants and Arx-1^+/-^ controls (Fig. 1E,F), we confirmed the migratory defects of the interneuron progenitors as reported by (Yoshihara et al., 2005) together with the absence of Arx-1 expression in the olfactory epithelium and in GnRH-1 neurons (Fig. 1G,H).

**Figure 1.**
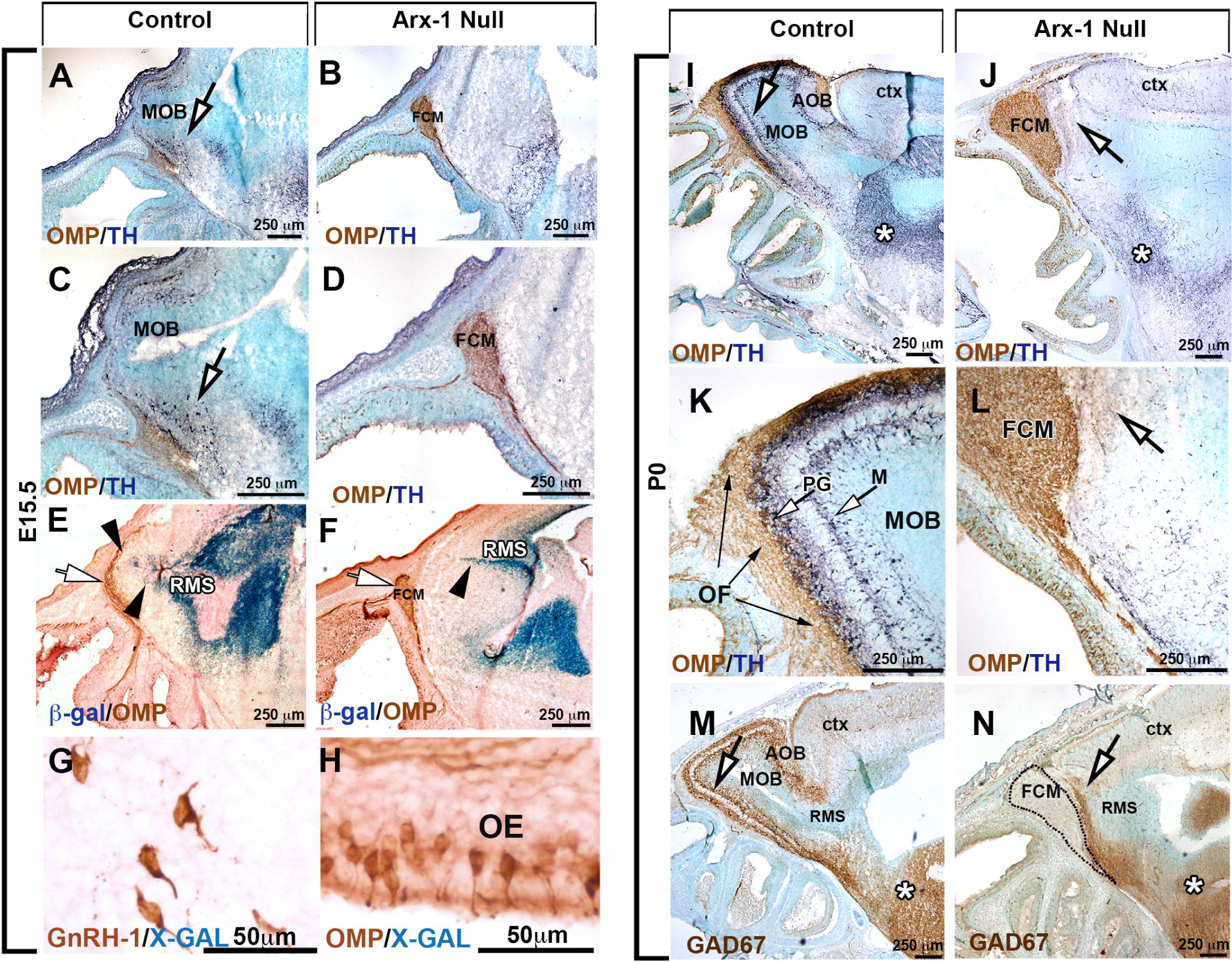
The olfactory bulbs fail to develop properly in Arx-1^null^ mutant mice. **A-D)** Immunohistochemistry against OMP (brown) and TH (dark blue) on control **(A,C)** and Arx-1^null^ **(B,D)** mouse at E15.5 shows detectable TH cells in the developing OB of control animals (arrow) but not in the mutants where the FCM stalls in front of the brain. OMP (brown) positive fibers were found projecting and targeting to the MOB of controls animals while they collapsed proximal to the brain as part of the FCM in Arx-1 mutants. **E-F)** OMP immunostaining and X-Gal staining on Arx-1 E15.5 Het control **(E)** and Arx-1^Null^ **(F)** In controls, Arx-1 was expressed in the rostral migratory stream and in cells invading the developing MOB (black arrows) innervated by the olfactory fibers (white arrow). In Arx-1^Null^ mutants the Arx-1+ cells fail to invade the developing MOB. **G-H)** X-Gal staining on E15.5 Arx-1 Het. Arx-1 expression (blue) is not found in **G)** GnRH-1 neurons nor in **H)** OMP+ olfactory neurons. P0 Controls **(I,K,M**) and Arx-1^null^ **(J,L,N).** Immunohistochemistry against **(I-L)** OMP (brown) and TH (dark blue) on control **(I,L)** and Arx-1^null^ mouse **(J,L)**. In the control, TH positive cells (white arrows) were found in the developing peri-glomerular (PG) and molecular layer (M), no TH+ cells were found in the Arx-1^null^ mutants **(L,N)** (arrow). **M,N)** GAD67 immunostaining on P0 **M)** control and **N)** Arx-1^null^ mutant. In control animals, GAD67 immunoreactivity was detected in the MOB (arrow), in mutant mice, GAD67+ cells accumulated ventrally and at the rostral end of the RMS (arrow). Comparable GAD67 pattern of immunoreactivity between controls and Arx-1^null^ was detected in the striatum (*).

### Aberrant olfactory development does not affect GnRH-1 migration to the basal forebrain

GnRH-1 neurons start to invade the brain region between E11.5 and E12.5 and complete their migration in about 5 days. We analyzed Arx-1^null^ mice and wild type controls at E13.5 and E15.5, which are stages in which the GnRH-1 neurons are still migrating, and at the completion of embryonic development, P0. Double immunolabeling against OMP and GnRH-1 (Fig. 2A-D) or Peripherin and GnRH-1 (Fig. 2I,J) revealed that despite the dramatic tangling of the olfactory and vomeronasal fibers observed in the Arx-1^null^ mutants, the GnRH-1ns were nonetheless able to bypass the tangle and access the brain (Fig. 2B,D,J). In both controls and Arx-1^null^ mutants, the GnRH-1ns were seen migrating to the brain along OMP-negative fibers (Fig. 2C,D).

**Figure 2.**
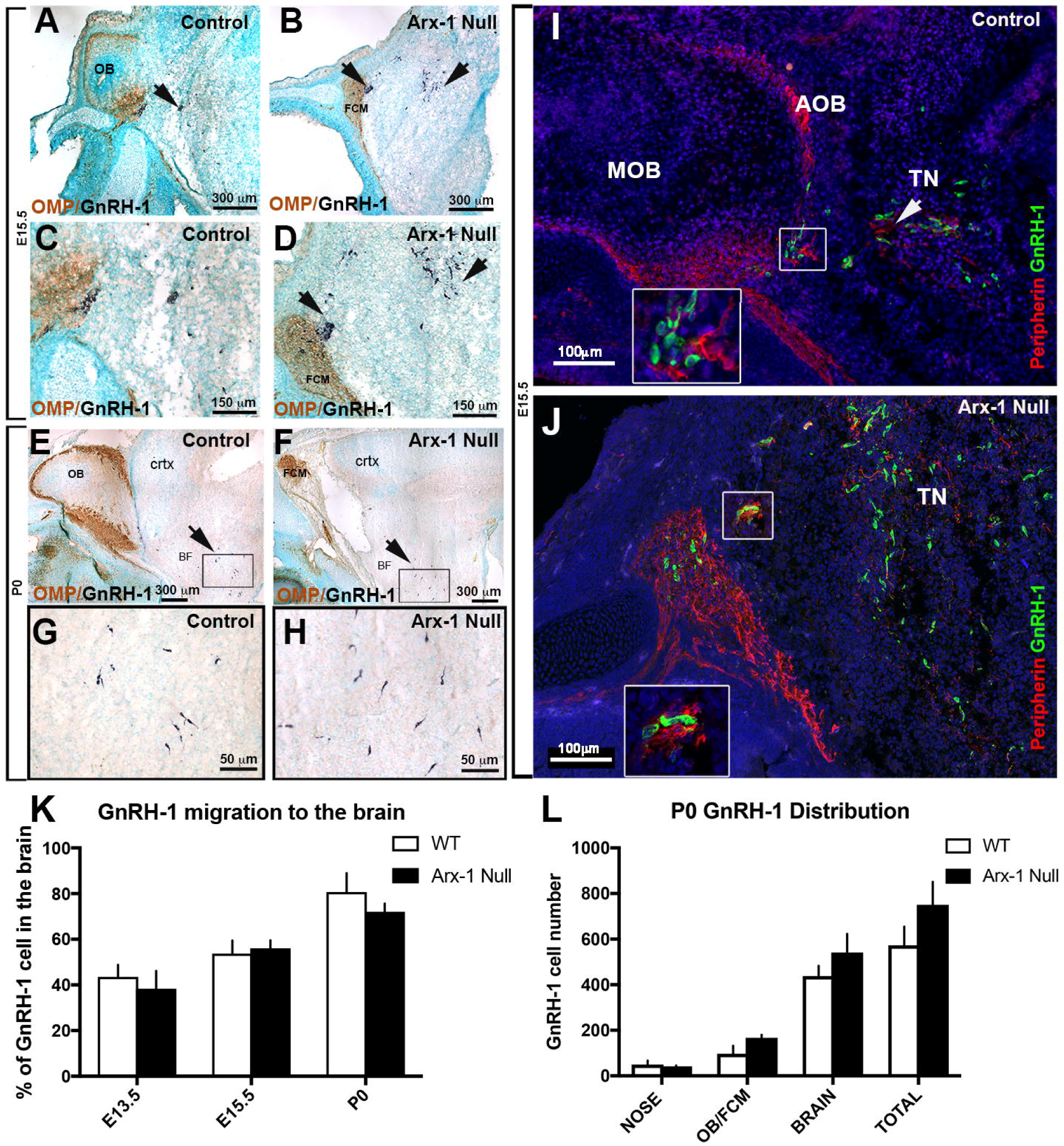
GnRH-1 neuron migration is not altered in Arx-1^null^ mutants. **A-H)** Immunohistochemistry against OMP (brown) and GnRH-1 (blue) on control **(A,C,E,G)** and Arx-1^null^ **(B,D,F,H)** mouse at E15.5 **(A-D)** and P0 **(E-H)**. In both controls and Arx-1^null^, the GnRH-1 neurons (dark blue) migrate to the basal forebrain on OMP negative fibers. In Arx-1^null^ mutants **(D)** the GnRH-1 neurons cross the axonal tangle of the FCM mass and migrate to the basal forebrain **(F,H)** as in control animals **(E,G)** (black arrows). **I,J)** E15.5, double immunofluorescence against Peripherin and GnRH-1. The GnRH-1 neurons migrate to the basal forebrain on Peripherin positive fibers of the TN on both control and Arx-1^null^ mice (white arrows). Enlargements illustrate the TN separating from the olfactory fibers that project to the MOB in control **(I)** or collapsed in the FCM in Arx-1^null^ mutants **(J)**. **K)** At the three analyzed stages a similar percentage of GnRH-1 neurons migrated to brain in controls and Arx-1^null^ mice. **L)** Graphs of GnRH-1ns cell counts at P0 in the nose, OB/FCM, brain and total. No statistical difference was seen in all areas between control and Arx-1^null^ (data ±SE, Unpaired t-test p>0.05)

Quantification of total GnRH-1 numbers at E13.5 (WT=715±87; KO=676±38; n=3), E15.5 (WT=727±76; KO=725±39; n=4) and P0 (WT=576±82; KO=661±99; n=3) indicated there were no differences between genotypes at any of the analyzed stages (data ±SE, unpaired t-test P>0.05).

To establish if the GnRH-1 neurons migrate in the brain at a different rate in Arx-1 mutants when compared to control animals, we quantified the distributions of GnRH-1ns between the nasal area and the brain at E13.5, E15.5, and P0 (Fig. 2K). This analysis revealed no difference among genotypes (Fig. 2K). Even in wild type mice, a subset of GnRH-1ns never reach the preoptic/hypothalamic area, but instead remain in the nasal area or form rings around the OBs (Casoni et al., 2016). Quantification of GnRH-1 neurons in the olfactory bulb/forebrain junction and brain (Fig. 2L), indicated that both at E15.5 (not shown) and P0, a similar number of GnRH-1ns remains in the FCM in Arx-1^null^ mutants compared with those found around the OBs in controls (Fig. 2L).

Performing a detailed mapping of GnRH-1 distribution in control and Arx-1^null^ mutants at P0, we observed comparable distribution of the GnRH-1 neurons, from the entry point, ventral to the olfactory bulbs, to the caudal hypothalamus, proximal to the median eminence (Fig. 3) to controls. However, the GnRH-1 cells in the Arx-1 mutants appeared to be less scattered along the dorso-ventral axis when compared to controls (Fig. 3C,D).

Thus, our analyses at these developmental stages argued that during embryonic development the GnRH-1ns migrate into the forebrain at comparable rate as in control mice, regardless of the severity of the olfactory bulb aplasia and the dramatic defects in olfactory axonal termination.

**Figure 3.**
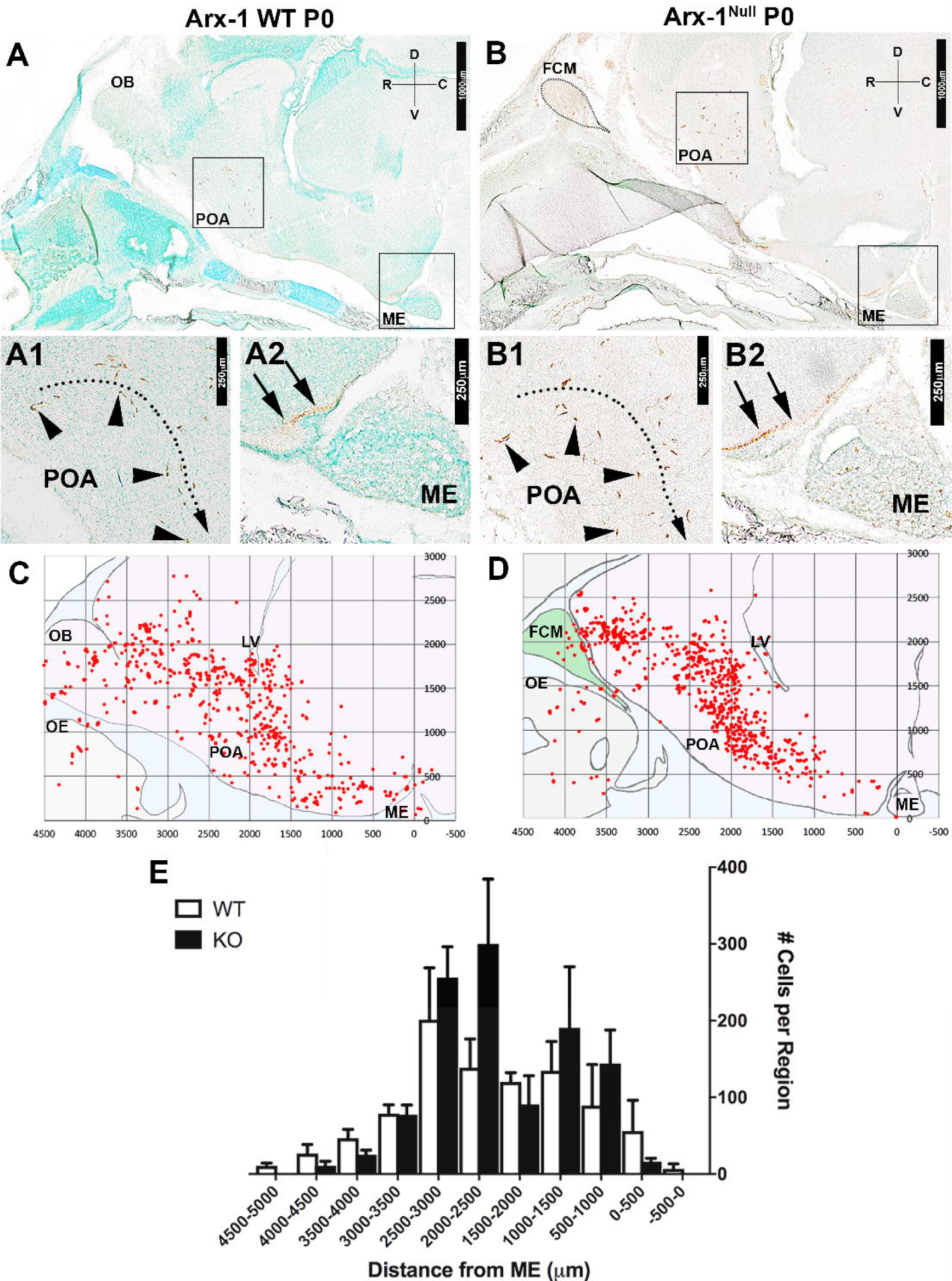
Aberrant formation of the olfactory system does not significantly affect the rostro-caudal distribution of GnRH-1ns in the brain. **A-B2)** Immunostaining against GnRH-1 on parasagittal sections on P0 Arx-1 WT (A-A2) and KO (B-B2) show similar distribution of A1,B1) GnRH-1ns cell bodies (arrowheads) in the POA and their A2,B2) projections (arrows) to the ME (D=dorsal, V=ventral, R=rostral, C=caudal). Mapping of the GnRH-1ns distribution in Arx-1 WT (n=3) and null (n=3) mice was performed on one series from each animal. C-D) Scatter plots illustrating GnRH-1ns distribution in WT (C) (n=3) and KO (n=3) (D), obtained by overlapping the GnRH-1ns coordinates from one series from each animal in reference to the median eminence (0,0 *μ*m), in along similar migratory paths. E) Histogram illustrating the average number of GnRH-1ns on the rostral-caudal axis in 500 *μ*m intervals. (data ±SE, Unpaired t-test p>0.05) (FCM=fibrocellular mass, LV=lateral ventricle, ME=median eminence, OB=olfactory bulb, POA=pre-optic area).

### The fibers upon which the GnRH-1 neurons access the brain are distinct from the olfactory fibers

Immunostaining for endogenous Peripherin in mice is usually used to highlight axons of cranial nerves, including those of the olfactory/vomeronasal and terminal nerves (TN/cranial nerve 0) (Wray et al., 1994; Yoshida et al., 1995; Schwanzel-Fukuda, 1999; Casoni et al., 2016). Thus, one of the major technical limitations in developmental studies of GnRH-1ns is the lack of molecular markers able to selectively label neuronal subpopulations in the nasal area. While analyzing the *hPRPH1-G* BAC transgenic line, which expresses EGFP under control of a human Peripherin gene promoter (McLenachan et al., 2008), we observed in the nasal area that expression of the hPeripherin:EGFP fusion protein was not fully consistent with that of the endogenous mouse gene. Whereas the endogenous mouse Peripherin protein was readily detectable by immunostaining for Peripherin on OSNs, VSNs, and on fibers forming the GnRH-1 migratory pathway (Fig. 4H,I,O), *hPRPH1-*EGFP expression was strong only in putative VSNs projecting to the accessory olfactory bulb (AOB) and in TN neurons and barely detectable in OSNs projecting to the main olfactory bulb (MOB) (Fig. 4A,B,J,L). Immunostaining against TAG-1, which was previously found to highlight neurons forming the GnRH-1 migratory scaffold (Yoshida et al., 1995; Casoni et al., 2016), and GnRH-1 confirmed that the EGFP+ fibers projecting to the basal forebrain were fibers of the presumptive TN (Fig. 4C-G).

**Figure 4.**
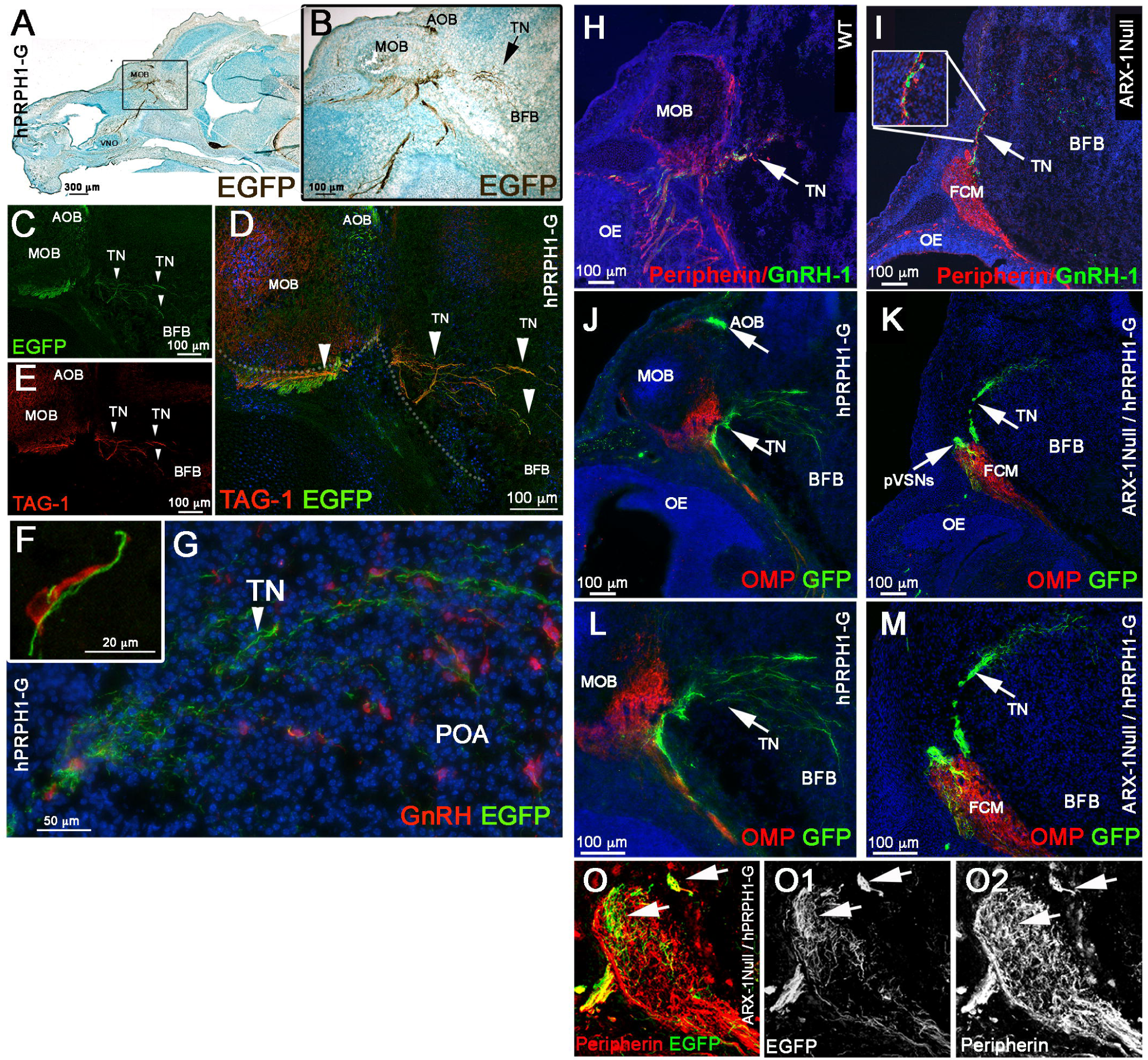
Expression of EGFP under the control of a human peripherin promoter distinguishes the terminal nerve (TN) from the olfactory/vomeronasal nerve. **A-G)** E15.5, parasagittal sections on hPRPH-1G. **A,B)** DAB immunostaining against EGFP showing strong EGFP expression in the AOB and TN invading the basal forebrain (BFB). **C-E)** Double immunofluorescence against TAG-1 and EGFP. both EGFP and TAG-1 were found to be strongly expressed in the TN fibers, while low TAG-1 expression was detected in putative vomeronasal fibers (VF) projecting to the AOB. **F,G)** E15.5, double immunofluorescence against EGFP and GnRH-1. GnRH-1ns (red) access the brain along Peripherin EGFP positive TN fibers. **(H-O) hPRPH1-G mice reveal that the TN is distinct form OSNs and bypasses the FCM in Arx-1^null^ mutants. H-I)** Peripherin/GnRH-1 double immunostaining on E15.5 WT **(H)** and Arx-1^null^ **(I)** shows GnRH-1 neurons entering the forebrain along the TN fibers positive for Peripherin (arrow). **I)** In the Arx-1^null^, GnRH-1 neurons access the brain along the Peripherin positive TN fibers that emerge from the Peripherin positive FCM. **J,L)** hPRPH1-G, E15.5, double immunostaining against OMP and EGFP reveals that the OMP+ olfactory sensory neurons projecting to the MOB are mainly negative for EGFP expression while the VSNs projecting to the AOB and the TN invading the basal forebrain (BFB) are positive for EGFP (arrows). **K,M)** double immunostaining against OMP and EGFp on hPRPH1-G /Arx1^null^, E15.5. The FCM is composed of OMP+ collapsed axons (red; tangled fibers) mainly negative for EGFP expression while the fibers of the TN, positive for <u>EGFP (arrows) are able to access the brain as in control animals (compare to K,M).</u> Putative (p) VSNs were found to be tangled as part of the FCM together with the OSNs strongly positive for OMP. **O-O2)** E15.5 hPRPH1-G /Arx1^null^ Double immunostaining against Peripherin and EGFP show hPRPH1-G is selective for pVSNs and the TN (arrows).

To validate that the GnRH-1ns follow the same migratory route in Arx-1^null^ mutants and controls, we exploited the stronger selective EGFP expression of the *hPRPH1-G* BAC transgenic line in VN and TN fibers (Fig. 4H-M). In line with what was observed after OMP/GnRH-1 immunolabeling (Fig. 2), immunolabeling against EGFP and OMP on *hPRPH1-G* mutant mouse sections (Fig. 4K,M) indicated that the fibers, upon which the GnRH-1 access the brain either did not express OMP or expressed it below immunodetectable levels. To follow selectively the trajectories of the putative TN in controls and Arx-1^null^ mice we generated *hPRPH1-G*^+/-^/ Arx-1^null^ embryos (Fig. 4K,M,O). Observations on these embryos revealed that the TN projections accessing the brain were positive for hPRP1-G expression, as was seen in control animals (Fig. 4J,L) while the hPRP1-G expressing vomeronasal sensory axons were tangled as part of the FCM (Fig. 4K,M,O).

### The TN fibers are distinct from apical and basal vomeronasal sensory neurons

Whereas some earlier researchers had proposed that GnRH-1ns reach the hypothalamus on a set of vomeronasal (VN) fibers that diverge from those that project to the accessory olfactory bulb (AOB) (Wray et al., 1989a; Yoshida et al., 1995), others argued instead that GnRH-1ns migrate along axons of the elusive terminal nerve (TN), which initially forms bundles with the vomeronasal axons until they diverge (Schwanzel-Fukuda and Pfaff, 1989; Vilensky, 2012; Zhao et al., 2013; Casoni et al., 2016). To resolve this discrepancy, we searched for genetic markers that could distinguish among the various subpopulations of axons resident in the nasal regions where GnRH-1ns migrate. The vomeronasal organ itself (VNO) is composed of apical and basal subpopulations of sensory neurons (VSNs). Both subpopulations project to the AOB, but they differ in their final targets, expression of adhesion molecules, Semaphorin receptors (Neuropilins), Slit receptors (Robos), VN receptors, G-protein subunits, and glycoproteins (Schwarting et al., 1994; Francia et al., 2014; Perez-Gomez et al., 2014). Whereas the apical neurons project to the anterior (a) portion of the AOB, the basal neurons project to the posterior (p) AOB. The neuronal branch upon which GnRH-1ns migrate shares transient expression of glycoproteins expressed by the basal, but not the apical VSNs (Yoshida et al., 1995). By analyzing microarray data obtained from vomeronasal tissue, data (not shown), we found that the G-protein coupled receptor GPR12, which is a high-affinity receptor for sphingosylphosphorylcholine (Ignatov et al., 2003), is expressed in the developing VNO (data not shown, *In Situ* available at Eurexpress.org). We next tested whether an EGFP-tagged version of GPR12 could be used to selectively label the vomeronasal axons, using the GENSAT GPR12-EGFP (Tg(Gpr12-EGFP) LD58Gsat) BAC transgenic mouse line. Analysis of these mice (Fig. 5) indicated that indeed the vomeronasal neurons, along with a few, sparse olfactory and microvillar neurons in the main olfactory epithelium (data not shown), expressed GPR12 from embryonic to postnatal stages. Immunostaining with two markers that can distinguish apical from basal VSNs, G*α*o (not shown) and G*α*i2 (Chamero et al., 2012), revealed that GPR12-EGFP was expressed in both the apical and basal VSNs, which respectively project to the anterior (a) and posterior (p) portions of the AOB (Fig. 5A-C). By immunostaining sections from these mice at E15.5 (n=3; Fig. 5D-F) for GnRH-1, Peripherin, and NRP2, we found that fibers of the TN were genetically distinct from these axons. Moreover, migrating GnRH-1ns were found to deviate from the GPR12-EGFP+ vomeronasal bundles projecting to the AOB (Fig. 5D) and to access the brain along Peripherin positive fibers negative for GPR12-EGFP, consistent with the presence of distinct TN fibers (Fig. 5F). Furthermore, the GPR12-EGFP expression allowed us to confirm that NRP2 expression was limited to the olfactory and apical VSN axons (Fig. 5E). In addition to the GnRH-1ns, we also observed sporadic migratory cells expressing GPR12 in their cell body migrating together with the GnRH-1ns (data not shown).

**Figure 5.**
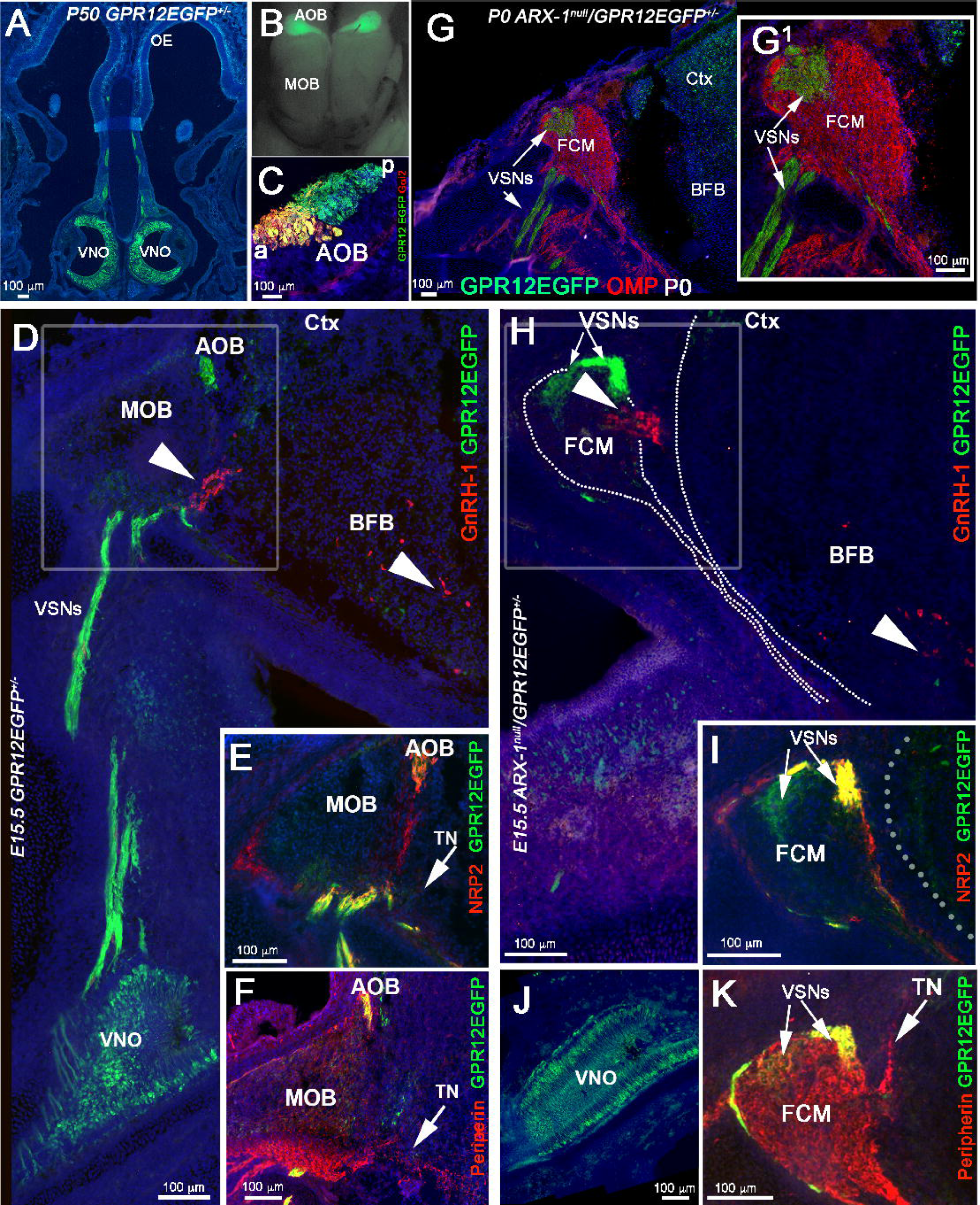
GPR12-EGFP BAC transgenics show that the TN is distinct from VSNs. **A-C)** Postnatal GPR12-EGFP. **A)** Coronal section, EGFP expression is limited to the VSNs and to sparse cells in the OE.**B)** Whole mount, EGFP is detectable in VSNs projecting to the AOB but not to the MOB. **C)** G*α*i2/EGFP staining on parasagittal section of the AOB, showing that the EGFP positive fibers project to both the anterior (a) and posterior (p) AOB. **D-F)** E15.5 GPR12-EGFP. **D)** Double immunostained against GnRH-1 and EGFP. The GnRH-1 neurons access the brain along GPR12-EGFP negative fibers (arrow heads). While GPR12-EGFP+ axons project from the VNO to the AOB. **E)** NRP2 (red) /EGFP (green) double staining shows NRP2 expression in GPR12-EGFP+ positive axonal bundles projecting to the AOB and in OSNs projecting to the MOB. **F)** Peripherin/EGFP double immunofluorescence, EGFP is expressed in the VSNs projecting to the AOB but not in the Peripherin+ TN (arrow). **G,G1)** P0, Arx-1^null^/GPR12-EGFP immunostained against OMP (red) and EGFP (green). The GPR12-EGFP positive vomeronasal fibers (VSNs) project toward the brain and collapse as part of the FCM. **H-K)** E15.5 Arx-1^null^/GPR12-EGFP. **H)** GnRH-1 (red) accessing the brain square, arrow in the BFB, on GPR12-EGFP negative fibers, the EGFP+ VSNs collapse as part of the FCM. **I)** NRP2 /EGFP double staining shows NRP2 expression in OSNs and in GPR12-EGFP+ positive VSNs in the FCM. **J)** EGFP expression in the VNO of E15.5 Arx-1^null^/GPR12-EGFP **K)** Peripherin/EGFP double immunofluorescence, Peripherin highlights the FCM and the TN emerging from the FCM, EGFP is expressed by the VSNs but not by the TN (arrows).

By mating Arx-1^null^ females with GPR12-EGFP males, we generated Arx-1^null^/GPR12-EGFP^+/-^ mice and GPR12-EGFP^+/-^ controls. Using these mice, we could selectively follow the trajectory of developing vomeronasal sensory fibers in the absence of proper olfactory bulb development by examining embryos at E15.5 and at birth (P0). In the Arx-1 mutants at P0 (Fig. 5G,G1), the VSNs axons formed a tangle within the FCM, surrounded by olfactory fibers. Analysis of Arx-1^null^/GPR12-EGFP^+/-^ embryos at E15.5 by immunostaining for EGFP in combination with GnRH-1, Peripherin, NRP2 (Fig. 5H-K) confirmed that the GnRH-1ns crossed the FCM and accessed the brain along neurons negative for GPR12EGFP. Collectively, these data showed that the axons of the TN are genetically distinct from vomeronasal axons, and are used by GnRH-1ns to access the brain.

### The GnRH-1 neurons and terminal nerve differ from the vomeronasal fibers for guidance receptors expression

The key regulators of olfactory axonal routing and targeting are the Class-3 Semaphorins, Neuropilin receptors (NRP-1 and NRP-2), Slit1, Slit2, and Roundabout (Robo) receptors (Renzi et al., 2000; Schwarting et al., 2000; Walz et al., 2002; Takeuchi et al., 2010; Cho et al., 2012). Slit and Sema3 proteins prevent olfactory fibers from invading the brain prior to OB formation (Renzi et al., 2000). Immunostaining against the Sema receptors NRP-1 and NRP-2 on hPRPH1-G^+/-^ mice showed that during GnRH-1 neuronal migration, NRP1 and NRP2 were strongly expressed by olfactory sensory neurons, projecting to the main olfactory (MOB) and by vomeronasal sensory neurons projecting to the accessory olfactory bulb, respectively (Fig. 6A-F). However, TN fibers, which are strongly EGFP+ in these mice, exhibited only weak NRP1 expression and no detectable NRP2 expression (Fig. 6A-F). Similarly, in Arx-1^null^/hPRPH1-G double mutants, we observed that NRP1 and NRP2 were expressed by the olfactory and vomeronasal fibers in the FCM, whereas the EGFP+ fibers of the TN expressed NRP1 only weakly, and no detectable NRP2 (Fig. 6G-H2).

**Figure 6.**
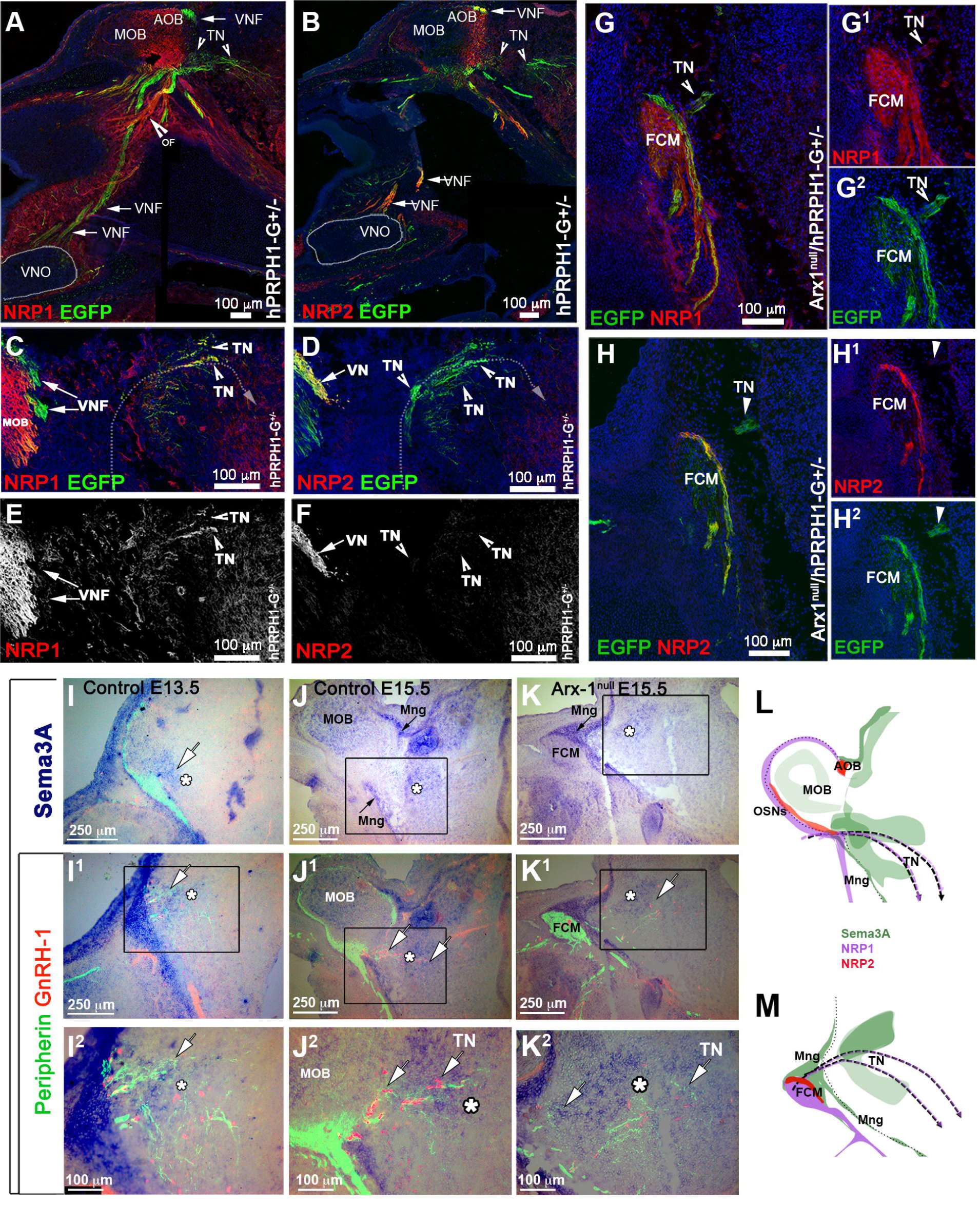
GnRH-1 neurons and the TN invade the brain proximal to a source of Sema3A. **A,C,E)** hPRPH1G^+/-^ E15.5, Immunostaining against NRP1 and EGFP. **A)** EGFP is expressed at high levels by putative VSNs projecting from the VNO to the AOB and by putative TN fibers accessing the brain ventral to the MOB. NRP1 immunoreactivity was not found along vomeronasal fibers (VNF) but in the nasal mesenchyme on the fibers of the OSNs neurons (OF) projecting to the MOB (yellow arrow). **C,E)** Enlargements showing the TN fibers accessing the brain express low levels of NRP1. **B,D,F)** hPRPH1G^+/-^ E15.5, Immunostaining against NRP2 and EGFP. NRP2 was strongly expressed the axonal fibers of the VSNs (VNF) by subsets of fibers of the OSNs neurons (OF) projecting to the MOB (arrows). **D,F)** Enlargements showing the TN fibers accessing the brain are negative or below detectability for NRP2 (notched arrows). **G-H2)** Arx1^null^/hPRPH1-G^+/-^ E15.5. **G-G2)** Immunostaining against NRP1 and EGFP reveals that while NRP1+ olfactory fibers are repelled from the developing telencephalon and collapse as part of the FCM the fibers of the TN, positive for EGFP (notched arrow) branch out of the FCM and project towards the brain. **H-H2)** IF against EGFP and NRP2, putative VN fibers positive for NRP2 and for EGFP were found to be part of the FCM while the fibers of the putative TN branch out of the FCM and project towards the brain. **I-K2)** ISH against Sema3A and IF against Peripherin and GnRH-1 on E13.5 **(I-I2)** and E15.5 **(J-K2)** controls **(I-J2)** and Arx-1^null^ **(J)** showing Sema3A expression if the forebrain (asterisk). In both controls and Arx-1^null^ mutants, strong Sema3A expression was found on the meninges (Mng, arrows) around the brain. In Arx-1^null^ mutants strong Sema3A was expressed on the meninges in contact with the FCM. **(K-K2)** IF against Peripherin and GnRH-1 on ISH against Sema3A reveals that the GnRH-1 neurons and the TN enter the brain in correspondence and in close proximity to a large source of Sema3a in Arx-1^null^ and controls. **I2,J2,K2)** Enlargements showing the TN fibers and GnRH-1ns accessing the brain through regions of Sema3a. **L,M)** Cartoons summarizing the trajectories of the Olfactory, vomeronasal and TN (dotted line) with respect to NRP1, NRP2 expression and sources of Sema3A in the brain and meninges (Mng).

To further understand the relationship between the aberrant olfactory/vomeronasal trajectories and successful GnRH-1 migration in Arx-1^null^ mutants, we performed *in situ hybridization* against the diffusible guidance cues Semaphorin 3A (Sema3A). By combining this digoxigenin-based in situ hybridization with double immunofluorescence for Peripherin and GnRH-1, we could follow the TN trajectory with respect to this guidance cue in the brain.

Analysis of WT animals at E13.5 and E15.5 consistently showed the TN and the GnRH-1ns invade the brain ventral and between the developing OBs in a region positive for Sema3A expression (Fig. 6I1-K2).

Consistent with observations in WT, analysis of Arx-1^null^ mutants at E15.5 showed that the FCM, which is mainly formed by NRP1+ fibers (Fig. 6G-G2) collapsed in close proximity with meninges positive for Sema3A expression (Fig. 6K,K1). However, the GnRH-1 neurons and TN were found to be able to penetrate the brain and to project towards and across sources of Sema3A (Fig. 6K1-2).

### The GnRH-1 neurons and terminal nerve respond differently from the olfactory and vomeronasal fibers to sources of Slit1 in the brain

Slit proteins play a pivotal role in repelling Robo1+ and Robo2+ olfactory and vomeronasal neurons and in preventing them from invading the brain (Renzi et al., 2000; Nguyen-Ba-Charvet et al., 2008). Consistent with this report, we observed, by in situ hybridization at E15.5, that both the olfactory and vomeronasal neurons expressed Robo2 (Fig. 7B1). Similarly, immunofluorescence confirmed detectable Robo2 expression in axons of the developing OSNs as well as in subsets of VSN projecting to the OB and tangled in the FCM of the Arx-1^Null^ mutants (Fig. 8H,I). Also, Robo1 expression was detected in putative olfactory ensheathing cells (OECs) (Aoki et al., 2013) surrounding the olfactory bulb in controls and surrounding the FCM in Arx-1^null^ mutants (Fig. 7A1-3, Fig. 8F,G). Consistent with a repellant role for Slit proteins in repelling Robo1+ and Robo2+ axons, in situ hybridization for Slit1 on controls and Arx-1^null^ mutants revealed strong Slit1 expression in the cortex as well as in the basal forebrain (Fig. 8 A-E3).

In sharp contrast to the OSNs and VSNs, neither the GnRH-1ns nor the terminal nerve expressed detectable levels of either Robo1 (Fig. 7A1-3, Fig. 8F,G) or Robo2 (Fig. 7B2-4).

**Figure 7.**
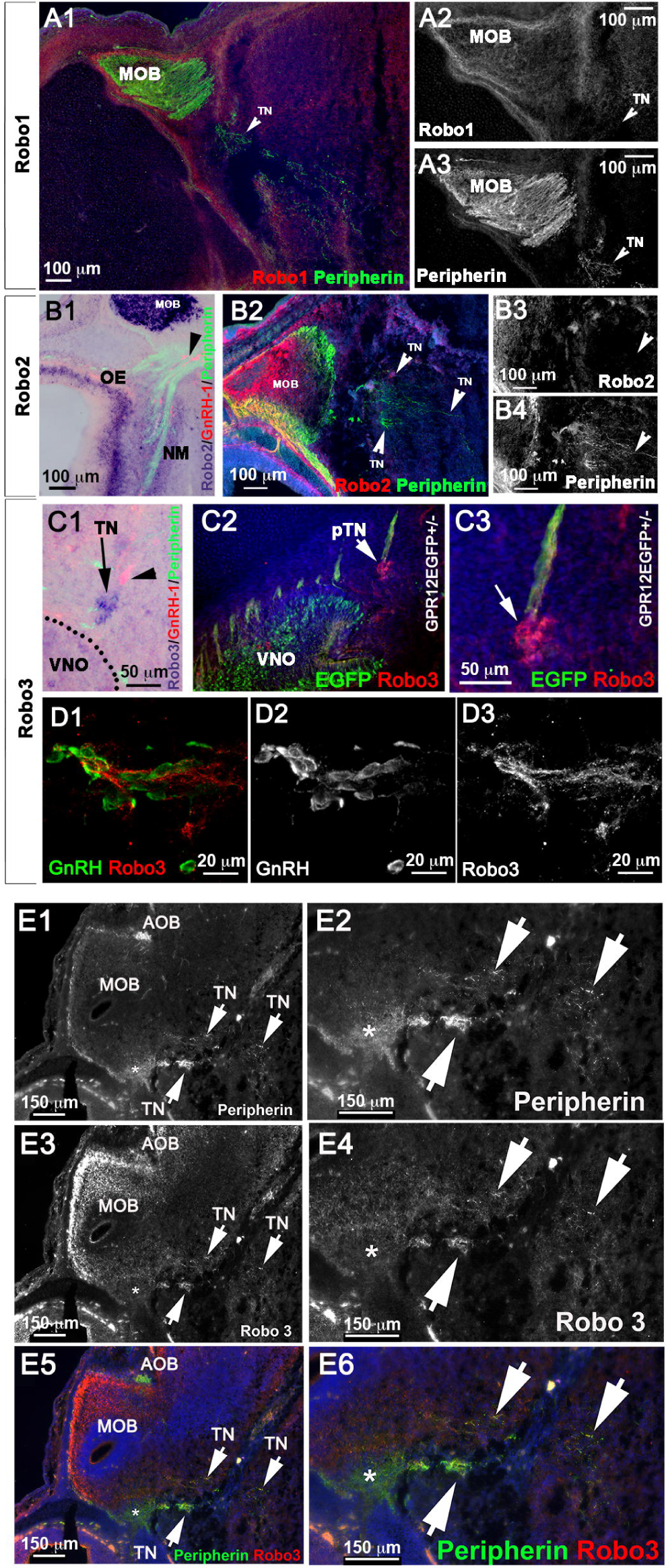
The TN is positive for Robo3 but not for Robo1 or Robo2. **A-B4)** E15.5 WT animal. A1-3) Double immunostaining against Robo1 and Peripherin show that the Peripherin+ TN is negative for Robo1 expression. B1) In-situ hybridization against Robo2 combined with immunofluorescence anti Peripherin and GnRH-1. Robo2 was detected in the vomeronasal neurons, olfactory epithelium (OE), nasal mesenchyme (NM), and in the olfactory bulb. No Robo2 signal was detected in GnRH-1 neurons. B2-4) Immunofluorescence against Robo2 and Peripherin shows lack of immunoreactivity for Robo2 in the TN accessing the brain. **C1)** E15.5 WT animal, in-situ hybridization against Robo3 combined with IF against GnRH-1 and Peripherin. Strong Robo3 mRNA expression was found in cells proximal to the VNO negative for GnRH-1 immunoreactivity (arrow). **C2-3)** E15.5 GPR12-EGFP immunostained for EGFP and Robo3 confirms strong Robo3 expression in cell bodies and fibers of neurons proximal to the VNO forming bundles with GPR12-EGFP+ VSNs (arrows in C2,3). Robo3 was found to be expressed at low levels in the vomeronasal organ and OE. Robo3+ cells proximal to the VNO negative for GnRH-1 and EGFP are indicated as putative cells of the terminal nerve (pTN). **D1-3)** E15.5 WT animal. GnRH-1 and Robo3 double immunofluorescence reveals migrating GnRH-1 neurons in contact with Robo3+ fibers. **E1-E6)** Double immunofluorescence against Robo3 and Peripherin in WT animals (E15.5). The fibers of the putative TN accessing the brain are positive for Robo3 and Peripherin immunoreactivity.

A third member of the Robo gene family of receptors, Robo3, does not bind Slit proteins, but various isoforms of Robo3 can silence Slit-mediated repulsion if they are co-expressed with Robo1 or Robo2 (Sabatier et al., 2004; Zelina et al., 2014). In-situ hybridization and immunohistochemistry against Robo3 revealed barely detectable levels of Robo3 expression in olfactory and vomeronasal neurons, but axons of the terminal nerve were strongly positive for Robo3 protein expression (Fig. 7E3-4). Furthermore, Robo3 mRNA was strongly expressed in cell bodies forming a ganglionic structure proximal to the VNO in the nasal area (Fig. 7C1). Robo3 immunostaining of GPR12-EGFP embryos confirmed that this staining was not in VSN cell bodies (Fig. 7C2-3). In the nasal region, Robo3 fibers comingled with GPR12-EGFP+ VSN fibers, and Peripherin immunostaining confirmed strong immunoreactivity for Robo3 on the fibers of the putative TN (Fig. 7C2-3,E1-6). Migrating GnRH-1ns were found along Robo3-expressing axons (Fig. 7D1-3).

To follow the trajectories of the OSN and VSNs with respect to sources of Slit in the brain at E13.5 and E15.5, we coupled in-situ hybridization against Slit1 with Peripherin and GnRH-1 immunofluorescence. At both stages, the OSN/VSNs contacted the brain in areas negative for Slit1 expression, which correspond to the putative olfactory bulb primordia (Fig. 8A1-2,C,D2). However, in Arx-1^null^ mutants, where the OBs fail to form properly, we found Slit1 to be expressed in a continuum throughout the cortex and basal forebrain (Fig. 8 B1-2,C,E1-2). Whereas the OSNs and VSNs positive for Robo1 and Robo2 (Fig, 8F-I) collapse proximal to sources of Slit1 (Nguyen-Ba-Charvet et al., 2008) (Fig 8B1-3,E2,C), the TN and GnRH-1 neurons appeared to be able to invade the brain and to migrate across sources of Slit1 (Fig 8 A3,B3, D3,E3). Collectively, these results (see summary in Fig. 8.C and J) suggest that GnRH-1ns migrate from the nose to the brain along axons of the TN. Our experiments indicate that routing of the axonal projections of the TN is defined by signaling mechanisms distinct from those controlling olfactory and vomeronasal targeting to the main and accessory olfactory bulbs.

**Figure 8.**
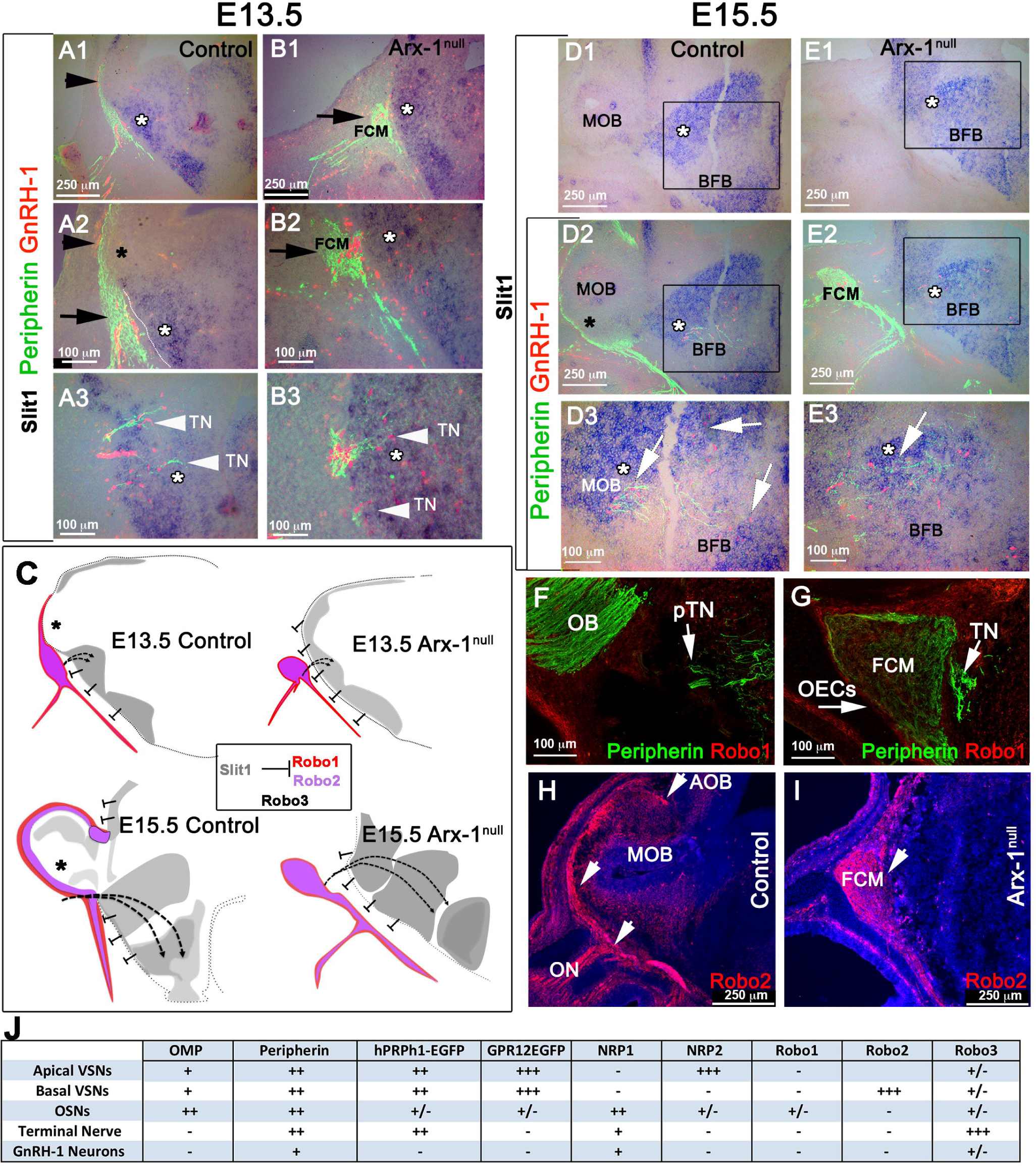
TN and GnRH-1 neurons invade the brain in areas of Slit1 expression. **A-B3)** In-situ hybridization against Slit1 (blue) combined with immunofluorescence against Peripherin and GnRH-1 in E13.5 WT **(A1-A3)** and Arx-1^null^ mutants **(B1-B3)**. **A1,A2)** Slit1 is strongly expressed in the cortex and basal forebrain (white asterisk) however the Peripherin positive ON and VSNs (black arrowhead) contact the brain the area negative for Slit1 (black asterisk) where the OB will form. **A3)** GnRH-1 neurons (red) invade the brain along Peripherin positive TN crossing a large source of Slit1. **B-B3)** In the Arx-1^Null^ the entire rostral border of the brain expresses Slit-1. As the OB primordium is failing to form, the olfactory and vomeronasal fibers did not access the brain and instead collapse, forming the FCM (black arrowhead) facing areas of Slit1 expression (white arrow). **B2-B3)** The GnRH-1 neurons (red) cross the FCM and start to penetrate the brain (white arrows) in Slit1 expression (white asterisk) areas as in control animals. **C)** Cartoon model illustrating the relationship between Robo1, Robo2, Robo3 and Slit1 in controls and Arx-1^null^ mutants during development of the olfactory and GnRH-1 system, The TN trajectories have been indicated with dotted line. Slit free areas indicated by black asterisks. **D-E3)**, In-situ hybridization against Slit1 (blue) combined with immunofluorescence against Peripherin and GnRH-1 in E15.5 WT (D-D3) and Arx-1^null^ mutants (E-E3). Slit1 was found to be strongly expressed (white asterisk) in the cortex and basal forebrain. In control animals, the olfactory and VSNs Peripherin+ fibers were found to project to the OB, which is mainly negative for Slit1 (black asterisk) while in the KOs the olfactory and vomeronasal fibers, did not access the brain strongly positive for Slit1. In both controls **(D2-3)** and Arx-1^null^ **(E2-3)** GnRH-1 neurons and Peripherin positive TN fibers (white arrows) were found to be able to access the brain crossing large sources of Slit1 (white asterisk). **F)** WT E15.5, Robo1/Peripherin immunofluorescence shows lack of Robo1 expression in the TN accessing the brain (arrows). **G)** E15.5 Arx-1^null^. Robo1 immunofluorescence was detected on the olfactory ensheathing cells (OECS) surrounding and within the FCM (arrowhead), no Robo1 immunoreactivity was found in the TN (arrows). **H)** WT E15.5, Robo2 immunofluorescence shows Robo2 expression in the olfactory fibers projections to the MOB and VSNs projections to the posterior AOB. **I)** Arx-1^null^ (E15.5) Robo2 immunofluorescence shows broad Robo2 expression in the axons of the FCM facing the source of Slit1 (D2). **J)** Summary of the molecular differences found between olfactory, vomeronasal, GnRH-1 and neurons of the putative terminal nerve.

## Discussion

After the initial description of Kallmann syndrome (Kallmann FJ, 1944) and subsequent discovery that GnRH-1ns migrate from the nose to the brain (Schwanzel-Fukuda and Pfaff, 1989; Wray et al., 1989a; Wray et al., 1989b), a link between olfactory development and GnRH-1 migration was proposed, investigated, and accepted (Cariboni et al., 2007; Toba et al., 2008; Wray, 2010; Lewkowitz-Shpuntoff et al., 2012). This found further support in: 1) evidence indicating that the GnRH-1ns originate from the olfactory placode; 2) a correlation between HH and mutations in genes affecting innervation of the olfactory bulbs; 3) analyses of mice carrying mutation in guidance signals that can affect olfactory development (Wierman et al., 2011; Forni and Wray, 2015).

However, the incomplete penetrance of anosmia and HH in families carrying Kallmann syndrome (de Roux, 2005; Pitteloud et al., 2005; Trarbach et al., 2006; Karstensen and Tommerup, 2012; Moya-Plana et al., 2013) led us to question whether this link was truly causal. We thus analyzed GnRH-1 development in the Arx-1^null^ model, where the loss of the Arx-1 gene by precursors of OB interneurons severely compromises olfactory bulb development. The normally developing telencephalon releases repulsive cues to prevent the penetration of olfactory fibers, thereby directing them to the OBs (Renzi et al., 2000; Cloutier et al., 2002; Nguyen-Ba-Charvet et al., 2008). Thus, in Arx-1^null^ mutants as in other mouse models of arrhinencephaly, the absence of OBs forces the olfactory and vomeronasal sensory fibers to form axonal tangles where the OBs should be (Balmer and LaMantia, 2004; Imai et al., 2009).

Despite the brain defects, aberrant olfactory bulb formation, and the extreme misrouting of olfactory/vomeronasal axons, the GnRH-1 neuron’s migratory rate as well as their ability to reach the preoptic/hypothalamic areas was not obviously altered in Arx-1 mutants. Arx-1 loss affects normal development of the brain (Friocourt et al., 2006; Simonet et al., 2015), therefore, some differences in how cells scatter in the brain could reflect brain abnormalities (Fig.3). These results indicate that correct targeting of olfactory and vomeronasal axons to the OB plays no direct role in defining the rate, and routing of GnRH-1ns migration into the forebrain. Instead, GnRH-1ns appeared to migrate along the axons of the TN to access the brain.

By exploiting hPRPH-1G and GPR12-EGFP BAC transgenics we revealed a distinction between the TN and the olfactory and vomeronasal sensory neurons. The GPR12-EGFP BAC transgenic was found to be selectively expressed by VSNs, few olfactory neurons, but not by the TN neurons. Though we cannot exclude that some of the cell bodies of the TN might be within the developing VNO, our data points to a distinct genetic identity for this nerve from the VSNs (Yoshida et al., 1995).

A small number of GnRH-1ns was found to fail to enter the brain in the FCM of Arx-1^null^ mutants. However, even in normal animals, a similar number of GnRH-1ns migrate on fibers projecting to the OB (Casoni et al., 2016). This suggests the existence of a subpopulation of GnRH-1ns that invariably migrates to the OB along specific neurons that must genetically differ from the majority that migrate along projections to the hypothalamus. If the GnRH-1ns that migrate to the OBs (Casoni et al., 2016) play active roles in olfaction is a possibility that should be further investigated.

Strengthening the idea that the TN and not the olfactory/vomeronasal fibers provides the scaffold for GnRH-1ns migration, we showed that these different subpopulations of axons follow different guidance cues. Targeting of axons in general is defined by a complex interplay of attractive and repulsive signals, which in the case of olfactory fibers, are released by the olfactory/vomeronasal epithelia, nasal mesenchyme, and the olfactory bulb (Schwarting et al., 2000; Cloutier et al., 2002; Cloutier et al., 2004; Cho et al., 2007; Prince et al., 2009; Cho et al., 2012; Brignall and Cloutier, 2015). An array of GnRH-1 migratory defects, with varying severity, occurs in genetically modified animal models where axonal misrouting and/or defasciculation of olfactory neurons also occurs (Matsumoto et al., 2006; Cariboni et al., 2011; Hernandez-Miranda et al., 2011; Messina et al., 2011; Cariboni et al., 2012; Barraud et al., 2013; Pingault et al., 2013; Cariboni et al., 2015; Tillo et al., 2015). However, discriminating between cell-autonomous and secondary effects of mutations of the genes linked to KS in humans (*e.g.,* prokineticin-2, prokineticin receptor-2, Fgf8, Fgf8-Receptor-1, Semaphorin3A, Semaphorin7A, Sox10, and CHD7) is made difficult by the broad number of tissue/cell types affected (Hanchate et al., 2012; Lewkowitz-Shpuntoff et al., 2012). Semaphorin 3A, NRP1, NRP2, Slit proteins, and the receptor Robo3 have all been previously implicated in guiding olfactory axons and GnRH-1ns (Cariboni et al., 2011; Cariboni et al., 2012).

By performing in situ hybridization against Sema3A and Slit1 on control and Arx-1^null^ mice we confirmed that the forebrain is a large source of repellent molecules, which in concert, might prevent the olfactory and vomeronasal fibers, but not the TN nor the GnRH-1ns, from invading the brain (Renzi et al., 2000; Nguyen-Ba-Charvet et al., 2008). As shown previously, we observed that the Sema3 receptors, NRP1 and NRP2, are expressed by olfactory and vomeronasal neurons projecting to different regions of the MOB and AOB (Cloutier et al., 2002; Walz et al., 2002; Cloutier et al., 2004; Taku et al., 2016); however we could only detect NRP1 but not NRP2 immuno-reactivity on GnRH-1ns and TN axons (Cariboni et al., 2011; Hanchate et al., 2012; Giacobini and Prevot, 2013; Giacobini, 2015). By performing in situ hybridization against Sema3A in combination with immunofluorescence for GnRH-1 and Peripherin, we observed that the TN and GnRH-1 invade the brain crossing a source of Sema3A.

Loss of Sema3A expression compromises both olfactory/vomeronasal axon trajectories as well as those of the TN, along with GnRH-1ns migration. The effects of class 3 Semaphorins on growth cone trajectories varies according to Sema3 concentration, NRP expression levels, and levels of cyclic nucleotides, which can selectively favor growth cone attraction vs. repulsion (Song et al., 1998; Pond et al., 2002; Manns et al., 2012). In Arx-1 mutants, NRP1+ and NRP2+ olfactory and VSNs axons collapsed proximal to the brain as part of the FCM (Fig. 6), whereas in both wild type and Arx-1^null^ mutants, the TN projected towards a source of Sema3A. These observations, together with the phenotype of Sema3A KO, where TN and GnRH-1 fail to invade the brain (Cariboni et al., 2011; Giacobini and Prevot, 2013), suggest that Sema3A may prevent olfactory and vomeronasal fibers from invading the brain while attracting GnRH-1ns and TN fibers.

Robo1,2 receptors cause axonal collapse in response to Slit proteins, whereas Robo3, depending on the isoform, can silence Slit repulsion (Chen et al., 2008). In line with previous studies, we found that both olfactory neurons and subsets of vomeronasal neurons express Robo2, along with low levels of Robo1 and Robo3. In mice lacking Robo1,2 the olfactory fibers invade the forebrain, following a route similar to that followed by the TN (Nguyen-Ba-Charvet et al., 2008). However, despite significant olfactory defects, Robo1 and Robo2 double mutants have no GnRH-1 migratory defects (Cariboni et al., 2012). Also, our data on WT and Arx-1^null^ mutants showed that the cortex and basal forebrain are large sources of Slit1, and that the GnRH-1ns, in contrast to the olfactory and vomeronasal axons, cross sources of Slit proteins in the forebrain (Fig. 8). In line with these observations, TN and GnRH-1ns (data not shown) are negative for Robo1 and Robo2 expression, but positive for Robo3. A previous study described defects in GnRH-1 migration in Robo3^null^ animals, which was proposed to be in response to Robo3 binding Slit2 (Cariboni et al., 2012) and would seem to contradict our findings. However, Robo3 is now known to bind NELL2 and not Slit proteins (Camurri et al., 2005; Zelina et al., 2014; Jaworski et al., 2015). Therefore, defective GnRH-1 migration in Robo3 mutants could result from the inability of TN and GnRH-1ns to respond to NELL2 mediated guidance.

Our conclusion that the TN, and not the olfactory/vomeronasal sensory neurons, provides the scaffold for GnRH-1ns migration, is supported by comparative phylogenetic studies. For example, although the VNO is absent in birds, amphibians, and fish (Smith and Bhatnagar, 2000; Dulac and Torello, 2003), GnRH-1ns and the TN connecting the nose to the brain are present in these species (Demski and Schwanzel-Fukuda, 1987; Ridgway et al., 1987; Muske and Moore, 1988; Mousley et al., 2006; Zhao et al., 2013). Moreover, the TN provides the only connection between the nasal area and the brain in toothed whales (Odontoceti), animals in which the GnRH-1ns reach the brain but the VNO, OE, and OBs fail to form (Buhl and Oelschlager, 1986; Ridgway et al., 1987).

Our work has shown, for the first time, that the invasion of the brain by migrating GnRH-1ns is independent from the correct targeting of the olfactory and vomeronasal neurons to the OBs. Although an impaired sense of smell, absence or reduction of olfactory bulb volume are all common diagnostic parameters for Kallmann Syndrome, our work suggests that neither defects in olfactory bulb development nor aberrant olfactory/vomeronasal axonal routing are sufficient to prevent the migration of GnRH-1ns into the basal forebrain. The pathophysiological overlap between Kallmann syndrome and normosmic IHH in humans (Lewkowitz-Shpuntoff et al., 2012) implies, that although the development of the TN/GnRH-1 system and the olfactory system may rely on partially overlapping guidance cues, they follow different molecular mechanisms. To reach a full understanding of the molecular mechanisms leading to KS and normosmic IHH, the community should now attempt to isolate and fully characterize the cells of the TN nerve in different animal systems and humans.

## AKNOWLEDGEMENTS

We thank Drs. Yoshio Goshima (Yokohoma City University School of Medicine, Yokohoma, Japan; Semaphorins 3A, 3C, 3F, NRP-1 and NRP-2); Jean-Francois Cloutier (Montreal Neurological Institute, Centre for Neuronal Survival, Montréal, Québec, Canada; Slits-1,2,3 as well as Robo-2, Robo-3). We thank Dr. A. Poulos for sharing his scanning microscope and helping us with the image acquisitions, Dr. D. Zuloga for sharing the GAD67 antibody. We thank Dr. J. Sasero (Murdoch Childrens Research Institute, Melbourne Australia) for sharing the hPRPH1-G mouse line, Dr. Ben Szaro (University at Albany) for the critical reading of the manuscript and Dr. Susan Wray (NINDS, NIH) for supporting some of the initial observations of this work (Intramural Research Program of the National Institutes of Health, NINDS NS002824-23 to SW). This work was supported by SUNY startup funds.

## REFERENCES

Aoki M, Takeuchi H, Nakashima A, Nishizumi H, Sakano H (2013) Possible roles of Robo1+ ensheathing cells in guiding dorsal-zone olfactory sensory neurons in mouse. Dev Neurobiol 73:828–840.

Balasubramanian R, Choi JH, Francescatto L, Willer J, Horton ER, Asimacopoulos EP, Stankovic KM, Plummer L, Buck CL, Quinton R, Nebesio TD, Mericq V, Merino PM, Meyer BF, Monies D, Gusella JF, Al Tassan N, Katsanis N, Crowley WF, Jr. (2014) Functionally compromised CHD7 alleles in patients with isolated GnRH deficiency. Proc Natl Acad Sci U S A 111:17953–17958.

Balmer CW, LaMantia AS (2004) Loss of Gli3 and Shh function disrupts olfactory axon trajectories. The Journal of comparative neurology 472:292–307.

Barraud P, St John JA, Stolt CC, Wegner M, Baker CV (2013) Olfactory ensheathing glia are required for embryonic olfactory axon targeting and the migration of gonadotropin-releasing hormone neurons. Biol Open 2:750–759.

Berghard A, Hagglund AC, Bohm S, Carlsson L (2012) Lhx2-dependent specification of olfactory sensory neurons is required for successful integration of olfactory, vomeronasal, and GnRH neurons. FASEB journal: official publication of the Federation of American Societies for Experimental Biology.

Bergman JE, Bosman EA, van Ravenswaaij-Arts CM, Steel KP (2010) Study of smell and reproductive organs in a mouse model for CHARGE syndrome. Eur J Hum Genet 18:171–177.

Bianco SD, Kaiser UB (2009) The genetic and molecular basis of idiopathic hypogonadotropic hypogonadism. Nat Rev Endocrinol 5:569–576.

Boehm U, Bouloux PM, Dattani MT, de Roux N, Dode C, Dunkel L, Dwyer AA, Giacobini P, Hardelin JP, Juul A, Maghnie M, Pitteloud N, Prevot V, Raivio T, Tena-Sempere M, Quinton R, Young J (2015) Expert consensus document: European Consensus Statement on congenital hypogonadotropic hypogonadism--pathogenesis, diagnosis and treatment. Nat Rev Endocrinol 11:547–564.

Brignall AC, Cloutier JF (2015) Neural map formation and sensory coding in the vomeronasal system. Cell Mol Life Sci 72:4697–4709.

Buhl EH, Oelschlager HA (1986) Ontogenetic development of the nervus terminalis in toothed whales. Evidence for its non-olfactory nature. Anat Embryol (Berl) 173:285–294.

Burmeister SS, Jarvis ED, Fernald RD (2005) Rapid behavioral and genomic responses to social opportunity. PLoS Biol 3:e363.

Camurri L, Mambetisaeva E, Davies D, Parnavelas J, Sundaresan V, Andrews W (2005) Evidence for the existence of two Robo3 isoforms with divergent biochemical properties. Mol Cell Neurosci 30:485–493.

Cariboni A, Maggi R, Parnavelas JG (2007) From nose to fertility: the long migratory journey of gonadotropin-releasing hormone neurons. Trends Neurosci 30:638–644.

Cariboni A, Davidson K, Rakic S, Maggi R, Parnavelas JG, Ruhrberg C (2011) Defective gonadotropin-releasing hormone neuron migration in mice lacking SEMA3A signalling through NRP1 and NRP2: implications for the aetiology of hypogonadotropic hypogonadism. Hum Mol Genet 20:336–344.

Cariboni A, Andrews WD, Memi F, Ypsilanti AR, Zelina P, Chedotal A, Parnavelas JG (2012) Slit2 and Robo3 modulate the migration of GnRH-secreting neurons. Development 139:3326–3331.

Cariboni A, Andre V, Chauvet S, Cassatella D, Davidson K, Caramello A, Fantin A, Bouloux P, Mann F, Ruhrberg C (2015) Dysfunctional SEMA3E signaling underlies gonadotropin-releasing hormone neuron deficiency in Kallmann syndrome. J Clin Invest 125:2413–2428.

Carleton A, Petreanu LT, Lansford R, Alvarez-Buylla A, Lledo PM (2003) Becoming a new neuron in the adult olfactory bulb. Nat Neurosci 6:507–518.

Casoni F, Malone SA, Belle M, Luzzati F, Collier F, Allet C, Hrabovszky E, Rasika S, Prevot V, Chedotal A, Giacobini P (2016) Development of the neurons controlling fertility in humans: new insights from 3D imaging and transparent fetal brains. Development 143:3969–3981.

Cattanach BM, Iddon CA, Charlton HM, Chiappa SA, Fink G (1977) Gonadotrophin-releasing hormone deficiency in a mutant mouse with hypogonadism. Nature 269:338–340.

Chamero P, Leinders-Zufall T, Zufall F (2012) From genes to social communication: molecular sensing by the vomeronasal organ. Trends Neurosci 35:597–606.

Chen Z, Gore BB, Long H, Ma L, Tessier-Lavigne M (2008) Alternative splicing of the Robo3 axon guidance receptor governs the midline switch from attraction to repulsion. Neuron 58:325–332.

Cho JH, Kam JW, Cloutier JF (2012) Slits and Robo-2 regulate the coalescence of subsets of olfactory sensory neuron axons within the ventral region of the olfactory bulb. Dev Biol 371:269–279.

Cho JH, Lepine M, Andrews W, Parnavelas J, Cloutier JF (2007) Requirement for Slit-1 and Robo-2 in zonal segregation of olfactory sensory neuron axons in the main olfactory bulb. J Neurosci 27:9094–9104.

Chung WC, Moyle SS, Tsai PS (2008) Fibroblast growth factor 8 signaling through fibroblast growth factor receptor 1 is required for the emergence of gonadotropin-releasing hormone neurons. Endocrinology 149:4997–5003.

Cloutier JF, Giger RJ, Koentges G, Dulac C, Kolodkin AL, Ginty DD (2002) Neuropilin-2 mediates axonal fasciculation, zonal segregation, but not axonal convergence, of primary accessory olfactory neurons. Neuron 33:877–892.

Cloutier JF, Sahay A, Chang EC, Tessier-Lavigne M, Dulac C, Kolodkin AL, Ginty DD (2004) Differential requirements for semaphorin 3F and Slit-1 in axonal targeting, fasciculation, and segregation of olfactory sensory neuron projections. J Neurosci 24:9087–9096.

de Roux N (2005) Isolated gonadotropic deficiency with and without anosmia: a developmental defect or a neuroendocrine regulation abnormality of the gonadotropic axis. Horm Res 64 Suppl 2:48–55.

Demski LS, Schwanzel-Fukuda M (1987) The terminal nerve (nervus terminalis): structure, function, and evolution. Introduction. Ann N Y Acad Sci 519:ix–xi.

Dode C, Hardelin JP (2010) Clinical genetics of Kallmann syndrome. Annales d’endocrinologie 71:149-157. Dulac C, Torello AT (2003) Molecular detection of pheromone signals in mammals: from genes to behaviour. Nat Rev Neurosci 4:551–562.

Ehrlich ME, Grillo M, Joh TH, Margolis FL, Baker H (1990) Transneuronal regulation of neuronal specific gene expression in the mouse olfactory bulb. Brain Res Mol Brain Res 7:115–122.

Forni PE, Wray S (2015) GnRH, anosmia and hypogonadotropic hypogonadism - Where are we? Front Neuroendocrinol 36C:165–177.

Forni PE, Bharti K, Flannery EM, Shimogori T, Wray S (2013) The indirect role of fibroblast growth factor-8 in defining neurogenic niches of the olfactory/GnRH systems. J Neurosci 33:19620–19634.

Francia S, Pifferi S, Menini A, Tirindelli R (2014) Vomeronasal Receptors and Signal Transduction in the Vomeronasal Organ of Mammals. In: Neurobiology of Chemical Communication (Mucignat-Caretta C, ed). Boca Raton (FL).

Frasnelli J, Schuster B, Hummel T (2007) Subjects with congenital anosmia have larger peripheral but similar central trigeminal responses. Cereb Cortex 17:370–377.

Friocourt G, Poirier K, Rakic S, Parnavelas JG, Chelly J (2006) The role of ARX in cortical development. Eur J Neurosci 23:869–876.

Ghadami M, Morovvati S, Majidzadeh AK, Damavandi E, Nishimura G, Kinoshita A, Pasalar P, Komatsu K, Najafi MT, Niikawa N, Yoshiura K (2004) Isolated congenital anosmia locus maps to 18p11.23-q12.2. Journal of medical genetics 41:299–303.

Giacobini P (2015) Shaping the Reproductive System: Role of Semaphorins in Gonadotropin-Releasing Hormone Development and Function. Neuroendocrinology 102:200–215.

Giacobini P, Prevot V (2013) Semaphorins in the development, homeostasis and disease of hormone systems. Semin Cell Dev Biol 24:190–198.

Hanchate NK et al. (2012) SEMA3A, a gene involved in axonal pathfinding, is mutated in patients with Kallmann syndrome. PLoS genetics 8:e1002896.

Hardelin JP, Dode C (2008) The complex genetics of Kallmann syndrome: KAL1, FGFR1, FGF8, PROKR2, PROK2, et al. Sexual development: genetics, molecular biology, evolution, endocrinology, embryology, and pathology of sex determination and differentiation 2:181–193.

Hernandez-Miranda LR, Cariboni A, Faux C, Ruhrberg C, Cho JH, Cloutier JF, Eickholt BJ, Parnavelas JG, Andrews WD (2011) Robo1 regulates semaphorin signaling to guide the migration of cortical interneurons through the ventral forebrain. J Neurosci 31:6174–6187.

Hirata T, Nakazawa M, Yoshihara S, Miyachi H, Kitamura K, Yoshihara Y, Hibi M (2006) Zinc-finger gene Fez in the olfactory sensory neurons regulates development of the olfactory bulb non-cell-autonomously. Development 133:1433–1443.

Ignatov A, Lintzel J, Hermans-Borgmeyer I, Kreienkamp HJ, Joost P, Thomsen S, Methner A, Schaller HC (2003) Role of the G-protein-coupled receptor GPR12 as high-affinity receptor for sphingosylphosphorylcholine and its expression and function in brain development. J Neurosci 23:907–914.

Imai T, Yamazaki T, Kobayakawa R, Kobayakawa K, Abe T, Suzuki M, Sakano H (2009) Pre-target axon sorting establishes the neural map topography. Science 325:585–590.

Jaworski A, Tom I, Tong RK, Gildea HK, Koch AW, Gonzalez LC, Tessier-Lavigne M (2015) Operational redundancy in axon guidance through the multifunctional receptor Robo3 and its ligand NELL2. Science 350:961–965.

Kagoshima M, Ito T (2001) Diverse gene expression and function of semaphorins in developing lung: positive and negative regulatory roles of semaphorins in lung branching morphogenesis. Genes Cells 6:559–571.

Kallmann FJ BS (1944) The genetic aspects of primary eunuchoidism. J Ment Defic 48:203–236.

Karstensen HG, Tommerup N (2012) Isolated and syndromic forms of congenital anosmia. Clin Genet 81:210–215.

Kawano T, Margolis FL (1982) Transsynaptic regulation of olfactory bulb catecholamines in mice and rats. J Neurochem 39:342–348.

Kitamura K et al. (2002) Mutation of ARX causes abnormal development of forebrain and testes in mice and X-linked lissencephaly with abnormal genitalia in humans. Nat Genet 32:359–369.

Kiyokage E, Kobayashi K, Toida K (2017) Spatial distribution of synapses on tyrosine hydroxylase-expressing juxtaglomerular cells in the mouse olfactory glomerulus. J Comp Neurol 525:1059–1074.

Kosaka K, Aika Y, Toida K, Heizmann CW, Hunziker W, Jacobowitz DM, Nagatsu I, Streit P, Visser TJ, Kosaka T (1995) Chemically defined neuron groups and their subpopulations in the glomerular layer of the rat main olfactory bulb. Neurosci Res 23:73–88.

Leopold DA, Hornung DE, Schwob JE (1992) Congenital lack of olfactory ability. Ann Otol Rhinol Laryngol 101:229–236.

Lettieri A, Oleari R, Gimmelli J, V An, Cariboni A (2016) The role of semaphorin signaling in the etiology of hypogonadotropic hypogonadism. Minerva Endocrinol 41:266–278.

Levi G, Puche AC, Mantero S, Barbieri O, Trombino S, Paleari L, Egeo A, Merlo GR (2003) The Dlx5 homeodomain gene is essential for olfactory development and connectivity in the mouse. Molecular and cellular neurosciences 22:530–543.

Lewkowitz-Shpuntoff HM, Hughes VA, Plummer L, Au MG, Doty RL, Seminara SB, Chan YM, Pitteloud N, Crowley WF, Jr., Balasubramanian R (2012) Olfactory phenotypic spectrum in idiopathic hypogonadotropic hypogonadism: pathophysiological and genetic implications. J Clin Endocrinol Metab 97:E136–144.

Long JE, Garel S, Depew MJ, Tobet S, Rubenstein JL (2003) DLX5 regulates development of peripheral and central components of the olfactory system. The Journal of neuroscience: the official journal of the Society for Neuroscience 23:568–578.

Manns RP, Cook GM, Holt CE, Keynes RJ (2012) Differing semaphorin 3A concentrations trigger distinct signaling mechanisms in growth cone collapse. J Neurosci 32:8554–8559.

Maruska KP, Fernald RD (2011) Social regulation of gene expression in the hypothalamic-pituitary-gonadal axis. Physiology (Bethesda) 26:412–423.

Matsumoto S, Yamazaki C, Masumoto KH, Nagano M, Naito M, Soga T, Hiyama H, Matsumoto M, Takasaki J, Kamohara M, Matsuo A, Ishii H, Kobori M, Katoh M, Matsushime H, Furuichi K, Shigeyoshi Y (2006) Abnormal development of the olfactory bulb and reproductive system in mice lacking prokineticin receptor PKR2. Proceedings of the National Academy of Sciences of the United States of America 103:4140–4145.

McLenachan S, Goldshmit Y, Fowler KJ, Voullaire L, Holloway TP, Turnley AM, Ioannou PA, Sarsero JP (2008) Transgenic mice expressing the Peripherin-EGFP genomic reporter display intrinsic peripheral nervous system fluorescence. Transgenic Res 17:1103–1116.

Messina A, Giacobini P (2013) Semaphorin Signaling in the Development and Function of the Gonadotropin Hormone-Releasing Hormone System. Front Endocrinol (Lausanne) 4:133.

Messina A, Ferraris N, Wray S, Cagnoni G, Donohue DE, Casoni F, Kramer PR, Derijck AA, Adolfs Y, Fasolo A, Pasterkamp RJ, Giacobini P (2011) Dysregulation of Semaphorin7A/beta1-integrin signaling leads to defective GnRH-1 cell migration, abnormal gonadal development and altered fertility. Hum Mol Genet 20:4759–4774.

Mitchell AL, Dwyer A, Pitteloud N, Quinton R (2011) Genetic basis and variable phenotypic expression of Kallmann syndrome: towards a unifying theory. Trends Endocrinol Metab 22:249–258.

Mousley A, Polese G, Marks NJ, Eisthen HL (2006) Terminal nerve-derived neuropeptide y modulates physiological responses in the olfactory epithelium of hungry axolotls (Ambystoma mexicanum). J Neurosci 26:7707–7717.

Moya-Plana A, Villanueva C, Laccourreye O, Bonfils P, de Roux N (2013) PROKR2 and PROK2 mutations cause isolated congenital anosmia without gonadotropic deficiency. Eur J Endocrinol 168:31–37.

Mugnaini E, Oertel WH, Wouterlood FF (1984) Immunocytochemical localization of GABA neurons and dopamine neurons in the rat main and accessory olfactory bulbs. Neurosci Lett 47:221–226.

Muske LE, Moore FL (1988) The nervus terminalis in amphibians: anatomy, chemistry and relationship with the hypothalamic gonadotropin-releasing hormone system. Brain Behav Evol 32:141–150.

Ng KL, Li JD, Cheng MY, Leslie FM, Lee AG, Zhou QY (2005) Dependence of olfactory bulb neurogenesis on prokineticin 2 signaling. Science 308:1923–1927.

Nguyen-Ba-Charvet KT, Di Meglio T, Fouquet C, Chedotal A (2008) Robos and slits control the pathfinding and targeting of mouse olfactory sensory Axons. Journal of Neuroscience 28:4244–4249.

Perez-Gomez A, Stein B, Leinders-Zufall T, Chamero P (2014) Signaling mechanisms and behavioral function of the mouse basal vomeronasal neuroepithelium. Front Neuroanat 8:135.

Pingault V, Bodereau V, Baral V, Marcos S, Watanabe Y, Chaoui A, Fouveaut C, Leroy C, Verier-Mine O, Francannet C, Dupin-Deguine D, Archambeaud F, Kurtz FJ, Young J, Bertherat J, Marlin S, Goossens M, Hardelin JP, Dode C, Bondurand N (2013) Loss-of-Function Mutations in SOX10 Cause Kallmann Syndrome with Deafness. Am J Hum Genet 92:707–724.

Pitteloud N, Acierno JS, Jr., Meysing AU, Dwyer AA, Hayes FJ, Crowley WF, Jr. (2005) Reversible kallmann syndrome, delayed puberty, and isolated anosmia occurring in a single family with a mutation in the fibroblast growth factor receptor 1 gene. J Clin Endocrinol Metab 90:1317–1322.

Pitteloud N, Zhang C, Pignatelli D, Li JD, Raivio T, Cole LW, Plummer L, Jacobson-Dickman EE, Mellon PL, Zhou QY, Crowley WF, Jr. (2007) Loss-of-function mutation in the prokineticin 2 gene causes Kallmann syndrome and normosmic idiopathic hypogonadotropic hypogonadism. Proc Natl Acad Sci U S A 104:17447–17452.

Pitteloud N, Acierno JS, Jr., Meysing A, Eliseenkova AV, Ma J, Ibrahimi OA, Metzger DL, Hayes FJ, Dwyer AA, Hughes VA, Yialamas M, Hall JE, Grant E, Mohammadi M, Crowley WF, Jr. (2006) Mutations in fibroblast growth factor receptor 1 cause both Kallmann syndrome and normosmic idiopathic hypogonadotropic hypogonadism. Proceedings of the National Academy of Sciences of the United States of America 103:6281–6286.

Pond A, Roche FK, Letourneau PC (2002) Temporal regulation of neuropilin-1 expression and sensitivity to semaphorin 3A in NGF- and NT3-responsive chick sensory neurons. J Neurobiol 51:43–53.

Prince JE, Cho JH, Dumontier E, Andrews W, Cutforth T, Tessier-Lavigne M, Parnavelas J, Cloutier JF (2009) Robo-2 controls the segregation of a portion of basal vomeronasal sensory neuron axons to the posterior region of the accessory olfactory bulb. J Neurosci 29:14211–14222.

Renzi MJ, Wexler TL, Raper JA (2000) Olfactory sensory axons expressing a dominant-negative semaphorin receptor enter the CNS early and overshoot their target. Neuron 28:437–447.

Ridgway SH, Demski LS, Bullock TH, Schwanzel-Fukuda M (1987) The terminal nerve in odontocete cetaceans. Ann N Y Acad Sci 519:201–212.

Sabatier C, Plump AS, Le M, Brose K, Tamada A, Murakami F, Lee EY, Tessier-Lavigne M (2004) The divergent Robo family protein rig-1/Robo3 is a negative regulator of slit responsiveness required for midline crossing by commissural axons. Cell 117:157–169.

Schwanzel-Fukuda M (1999) Origin and migration of luteinizing hormone-releasing hormone neurons in mammals. Microsc Res Tech 44:2–10.

Schwanzel-Fukuda M, Pfaff DW (1989) Origin of luteinizing hormone-releasing hormone neurons. Nature 338:161–164.

Schwanzel-Fukuda M, Bick D, Pfaff DW (1989) Luteinizing hormone-releasing hormone (LHRH)-expressing cells do not migrate normally in an inherited hypogonadal (Kallmann) syndrome. Brain Res Mol Brain Res 6:311–326.

Schwarting GA, Drinkwater D, Crandall JE (1994) A unique neuronal glycolipid defines rostrocaudal compartmentalization in the accessory olfactory system of rats. Brain Res Dev Brain Res 78:191–200.

Schwarting GA, Kostek C, Ahmad N, Dibble C, Pays L, Puschel AW (2000) Semaphorin 3A is required for guidance of olfactory axons in mice. J Neurosci 20:7691–7697.

Simonet JC, Sunnen CN, Wu J, Golden JA, Marsh ED (2015) Conditional Loss of Arx From the Developing Dorsal Telencephalon Results in Behavioral Phenotypes Resembling Mild Human ARX Mutations. Cereb Cortex 25:2939–2950.

Smith TD, Bhatnagar KP (2000) The human vomeronasal organ. Part II: prenatal development. J Anat 197 Pt 3:421–436.

Song H, Ming G, He Z, Lehmann M, McKerracher L, Tessier-Lavigne M, Poo M (1998) Conversion of neuronal growth cone responses from repulsion to attraction by cyclic nucleotides. Science 281:1515–1518.

Stone DM, Wessel T, Joh TH, Baker H (1990) Decrease in tyrosine hydroxylase, but not aromatic L-amino acid decarboxylase, messenger RNA in rat olfactory bulb following neonatal, unilateral odor deprivation. Brain Res Mol Brain Res 8:291–300.

Takeuchi H, Inokuchi K, Aoki M, Suto F, Tsuboi A, Matsuda I, Suzuki M, Aiba A, Serizawa S, Yoshihara Y, Fujisawa H, Sakano H (2010) Sequential arrival and graded secretion of Sema3F by olfactory neuron axons specify map topography at the bulb. Cell 141:1056–1067.

Taku AA, Marcaccio CL, Ye W, Krause GJ, Raper JA (2016) Attractant and repellent cues cooperate in guiding a subset of olfactory sensory axons to a well-defined protoglomerular target. Development 143:123–132.

Teixeira L, Guimiot F, Dode C, Fallet-Bianco C, Millar RP, Delezoide AL, Hardelin JP (2010) Defective migration of neuroendocrine GnRH cells in human arrhinencephalic conditions. The Journal of clinical investigation 120:3668–3672.

Tillo M, Erskine L, Cariboni A, Fantin A, Joyce A, Denti L, Ruhrberg C (2015) VEGF189 binds NRP1 and is sufficient for VEGF/NRP1-dependent neuronal patterning in the developing brain. Development 142:314–319.

Toba Y, Tiong JD, Ma Q, Wray S (2008) CXCR4/SDF-1 system modulates development of GnRH-1 neurons and the olfactory system. Dev Neurobiol 68:487–503.

Trarbach EB, Costa EM, Versiani B, de Castro M, Baptista MT, Garmes HM, de Mendonca BB, Latronico AC (2006) Novel fibroblast growth factor receptor 1 mutations in patients with congenital hypogonadotropic hypogonadism with and without anosmia. J Clin Endocrinol Metab 91:4006–4012.

Vilensky JA (2012) The neglected cranial nerve: Nervus terminalis (cranial nerve N). Clin Anat.

Walz A, Rodriguez I, Mombaerts P (2002) Aberrant sensory innervation of the olfactory bulb in neuropilin-2 mutant mice. J Neurosci 22:4025–4035.

Wierman ME, Kiseljak-Vassiliades K, Tobet S (2011) Gonadotropin-releasing hormone (GnRH) neuron migration: initiation, maintenance and cessation as critical steps to ensure normal reproductive function. Front Neuroendocrinol 32:43–52.

Wray S (2010) From nose to brain: development of gonadotrophin-releasing hormone-1 neurones. J Neuroendocrinol 22:743–753.

Wray S, Grant P, Gainer H (1989a) Evidence that cells expressing luteinizing hormone-releasing hormone mRNA in the mouse are derived from progenitor cells in the olfactory placode. Proceedings of the National Academy of Sciences of the United States of America 86:8132–8136.

Wray S, Nieburgs A, Elkabes S (1989b) Spatiotemporal cell expression of luteinizing hormone-releasing hormone in the prenatal mouse: evidence for an embryonic origin in the olfactory placode. Brain Res Dev Brain Res 46:309–318.

Wray S, Key S, Qualls R, Fueshko SM (1994) A subset of peripherin positive olfactory axons delineates the luteinizing hormone releasing hormone neuronal migratory pathway in developing mouse. Developmental biology 166:349–354.

Yin W, Gore AC (2006) Neuroendocrine control of reproductive aging: roles of GnRH neurons. Reproduction 131:403–414.

Yoshida K, Tobet SA, Crandall JE, Jimenez TP, Schwarting GA (1995) The migration of luteinizing hormone-releasing hormone neurons in the developing rat is associated with a transient, caudal projection of the vomeronasal nerve. J Neurosci 15:7769–7777.

Yoshida M, Suda Y, Matsuo I, Miyamoto N, Takeda N, Kuratani S, Aizawa S (1997) Emx1 and Emx2 functions in development of dorsal telencephalon. Development 124:101–111.

Yoshihara S, Omichi K, Yanazawa M, Kitamura K, Yoshihara Y (2005) Arx homeobox gene is essential for development of mouse olfactory system. Development 132:751–762.

Yousem DM, Geckle RJ, Bilker W, McKeown DA, Doty RL (1996) MR evaluation of patients with congenital hyposmia or anosmia. AJR American journal of roentgenology 166:439–443.

Zelina P, Blockus H, Zagar Y, Peres A, Friocourt F, Wu Z, Rama N, Fouquet C, Hohenester E, Tessier-Lavigne M, Schweitzer J, Roest Crollius H, Chedotal A (2014) Signaling switch of the axon guidance receptor Robo3 during vertebrate evolution. Neuron 84:1258–1272.

Zhang G, Li J, Purkayastha S, Tang Y, Zhang H, Yin Y, Li B, Liu G, Cai D (2013) Hypothalamic programming of systemic ageing involving IKK-beta, NF-kappaB and GnRH. Nature 497:211–216.

Zhao Y, Lin MC, Farajzadeh M, Wayne NL (2013) Early development of the gonadotropin-releasing hormone neuronal network in transgenic zebrafish. Front Endocrinol (Lausanne) 4:107.

